# Genomic screening reveals UBA1 as a potent and druggable target in c-MYC-high TNBC models

**DOI:** 10.1101/2022.05.17.491908

**Authors:** Sheeba Jacob, Tia H. Turner, Jinyang Cai, Konstantinos V. Floros, Ann K. Yu, Colin M. Coon, Rishabh Khatri, Mohammad A. Alzubi, Charles T. Jakubik, Ynes M. Bouck, Madhavi Puchalapalli, Mayuri Shende, Mikhail G. Dozmorov, Sosipatros A. Boikos, Bin Hu, J. Chuck Harrell, Cyril H. Benes, Jennifer E. Koblinski, Carlotta Costa, Anthony C. Faber

**Author notes:** co-corresponding author **To whom correspondence should be addressed:**; Department of Oral and Craniofacial Molecular Biology, Virginia Commonwealth/Philips Institute, 521 N, 11th Street, Richmond, VA-23298-5045. Department of Oncology, Novartis Institute for Biomedical Research, Oncology, Basel, Switzerland (Carlotta Costa). **Author Contributions** S.J., C.C., C.H.B. and A.C.F. designed the study and analyzed the data. S.J., C.C., J.E.K., M.D., J.C.H., Y.M.B., C.H.B. and A.C.F. wrote the paper. C.H.B., J.C.H., C.C. and A.C.F. supervised the studies. S.J., T.H.T., J.Y., K.V.F., A.Y., C.C., R.K., and M.A.A. performed cell line and biochemical studies. S.J., M.A.A., B.H., and T.H.T. performed PDX studies. C.C., C.T.J. and M.D. performed CRISPR/Cas9 screen analyses. S.J., T.H.T., C.H.B., C.C., J.E.K. and A.C.F. were involved with the study design. All authors discussed the results and commented on the manuscript.

## Abstract

Triple negative breast cancer (TNBC) accounts for over 30% of all breast cancer-related deaths, despite accounting for only 10%–15% of total breast cancer cases. Targeted therapy development has largely stalled in TNBC, underlined by a lack of traditionally druggable addictions like receptor tyrosine kinases (RTKs). Here, through full genome CRISPR/Cas9 screening of TNBC models, we have uncovered the sensitivity of TNBCs to the depletion of the Ubiquitin-Like Modifier Activating Enzyme 1 (UBA1). Targeting UBA1 with the first in-class UBA1 inhibitor TAK-243 induced unresolvable ER-stress and activating transcription factor 4 (ATF4)-mediated upregulation of pro-apoptotic NOXA, leading to cell death. In five patient derived xenograft models (PDXs) of TNBC, TAK-243 therapy led to tumor inhibition or frank tumor regression. In an intracardiac metastatic model of TNBC, TAK-243 markedly reduced metastatic burden. Importantly, there was an order of magnitude greater sensitivity of TNBC lines to TAK-243 compared to normal tissue-derived cells. Lastly, c-MYC expression correlates with TAK-243 sensitivity and cooperates with TAK-243 to induce a stress response and cell death. We posit UBA1 is an important new target in TNBC expressing high levels of c-MYC.

**Significance:** Genomic screening of TNBC cell lines revealed broad sensitivity to depletion of the E1 ubiquitin enzyme, UBA1. Disrupting UBA1 with the first in-class inhibitor TAK-243 in TNBC models induces ER-stress through an ATF4-NOXA axis that is dependent on c-MYC, leading to apoptosis, in vitro and in vivo, primary tumor growth inhibition and metastatic inhibition.

## Introduction

TNBC accounts for up to 15% of BCs and is often lethal, in particular, to young, Black women (1). About 4 out of every 5 TNBC cases are basal-like, and they are extremely heterogeneous in terms of genomic alterations (2). Despite major advances in targeted therapy implementation in other BCs like estrogen receptor (ER)+ BC and *HER2*-amplified BC, outside of *BRCA* mutant TNBC where PARP inhibitors are approved (2-4), there is a critical need for new targeted therapies. Unfortunately, other than reoccurring *PTEN* deletions and a small number of *PIK3CA* mutant TNBCs, both potentially amenable to treatment with PI3K inhibitors, there are no reoccurring mutations that would suggest sensitivity to other clinically available kinase inhibitors.

The implementation of targeted therapies has been revolutionary in different cancer paradigms including breast cancer. ER inhibitors and HER2 inhibitors have both been implemented into breast cancer patient care, where nearly 90% of patients positive for one of these alterations now survive. TNBCs by definition are not positive for hormone receptors or excessive HER2 receptors. Re-occurring alterations are mostly limited to loss of PTEN (∼ 35 percent of cases), PIK3CA activating mutations (∼ 9 percent of cases) and loss of BRCA (19.5 percent of cases) (2). Even so, treatment with PI3K inhibitors have not been successful in TNBC, and PARP inhibitor therapy has not been as successful in TNBC (5) as it has in BRCA mutant ovarian cancer (6). Extreme heterogeneity may underlie some of the lack of consistency in response to both of these classes of inhibitors (7, 8). Despite this, TNBCs have many common characteristics– for instance, about 80% are derived from basal-like tissue, TNBCs in general possess strong immunosuppressive qualities including high PD-L1 expression (9), and TNBCs have a tendency to strike younger minority women, where it is largely more aggressive than other breast cancers (10). Altogether, the genomic and clinical evidence may suggest that implementation of targeted therapies will likely require an expansion of potential targets beyond the kinome.

## Results

### Genomic screening of TNBC reveals hits in the ubiquitin-proteasome system (UPS)

In order to capture the full scope of potential targets, we performed whole genome CRISPR/Cas9 screening, covering approximately 20,000 target genes with four single-guide (sg)RNAs targeting each gene, in two TNBC cell lines; BT-549 (derived from primary breast tumor) and MDA-MB-468 (derived from pleural effusion) (Fig. 1A). Following 21 days of selection, the percent of surviving cells with each sgRNA was calculated by STARS analyses (11) (top hits are listed in Sup. Table 1). Interestingly, the analyses uncovered several gene hits in the ubiquitin-proteasome system (UPS) including multiple ubiquitin specific peptidases (USPs) and proteasome subunits, as well as previously reported TNBC survival genes like MYC (Sup. Table 1 and 2) (12, 13). Among the highest-ranked genes in both cell lines was Ubiquitin-Like Modifier Activating Enzyme 1 (UBA1) (also known as UBE1) (Sup. Table 1 and 2). UBA1 is the E1 ubiquitin-activating enzyme and represents a critical node of control in the UPS (14, 15). Based on the sensitivity of both cell lines to UBA1 depletion, we expanded our evaluation to an additional 7 TNBC cell lines (HCC1806, HCC70, SUM149PT, MDA-MB-231, Hs578T, MDA-MB-436, HCC1937), (Sup Table 3) using 10 UBA1-targeting sgRNAs. All nine cell lines demonstrated sensitivity to depletion of UBA1 (Sup. Fig. S1), suggesting broad efficacy throughout diverse TNBC models. TAK-243 is a novel UBA1 inhibitor (15-17) currently in clinical evaluation. To assess the potential of TAK-243 in recapitulating the genetic screens results, fourteen TNBC cell lines and three normal-tissue derived cell lines of different origin were treated with TAK-243. We found a striking difference between the sensitivity of the TNBC cell lines and normal tissue-derived cells, suggesting that TAK-243 has the potential to specifically target TNBC cells (Fig. 1B).

**Figure 1.**
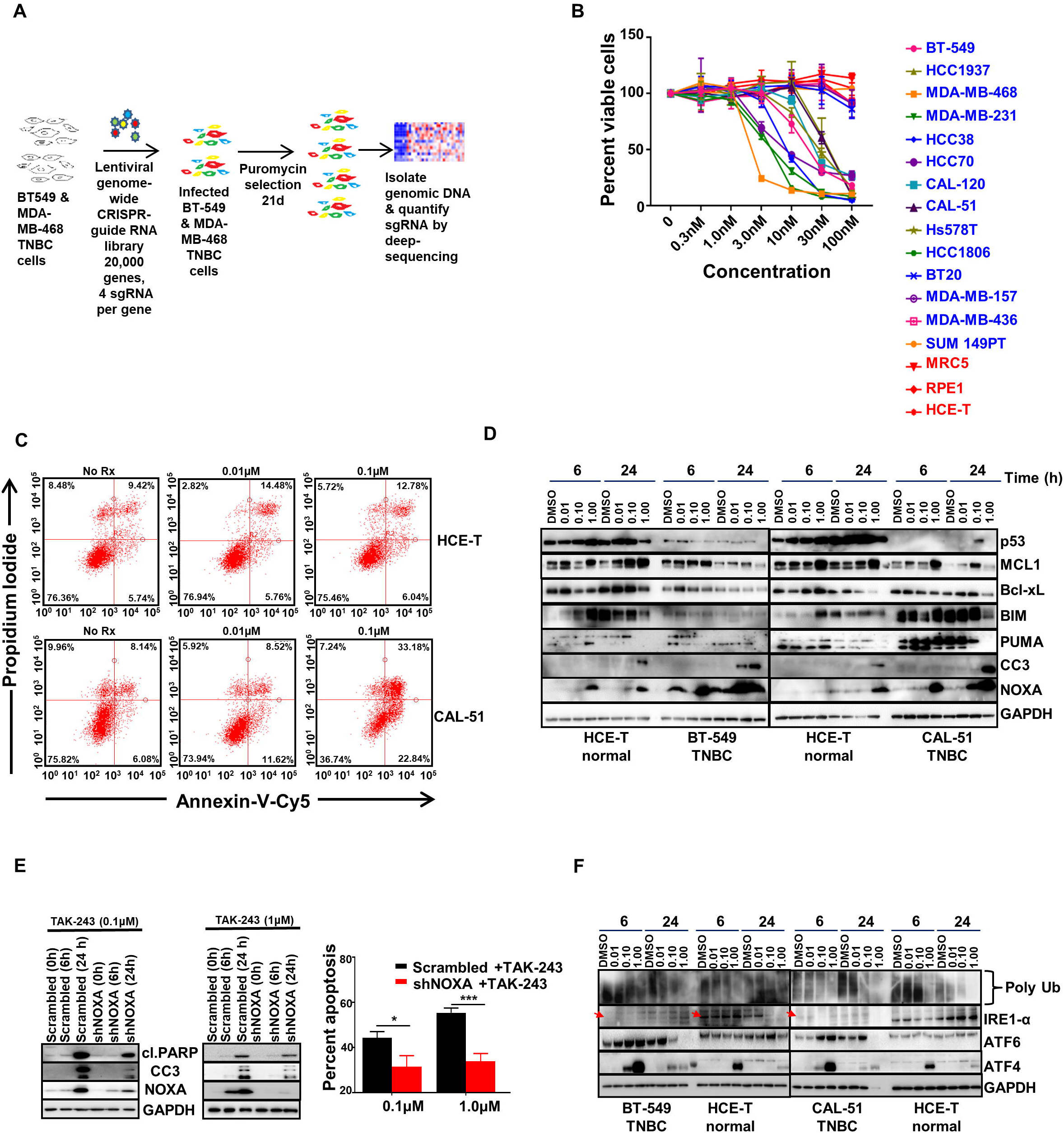
UBA1 is a target in TNBC. (A) Schema of TNBC genetic screen. Whole genome of the TNBC cell lines (BT-549 and MDA-MB-468) followed by UBA1 depletion in nine TNBC cell lines. (B) Graph represents percent viable cells assessed by CellTiter-Glo in TNBCs (blue) and normal tissue-derived MRC5, RPE1 and HCE-T cells (red) following 72 h treatment with TAK-243 at the indicated concentration. (C) FACS analysis showing annexin-V-Cy5 and propidium iodide staining in HCE-T and TNBC cells following 24h treatment with TAK-243 at indicated concentration (D) Western blot analysis showing dose response and time course of the effects of TAK-243 on apoptosis in TNBC (CAL-51 and BT-549) and normal HCE-T cells, as assessed by immunoblotting for p53, MCL-1, Bcl-xL, BIM, PUMA, cleaved Caspase 3 (CC3) and NOXA. GAPDH was used as a loading control. (E) Western blot analysis showing stable knockdown of NOXA. Reduced NOXA protects from TAK-243 toxicity in CAL-51 cells as assessed by immunoblotting for cleaved PARP, CC3 and NOXA. GAPDH was used as a loading control (*left* panel). Right panel indicates percent apoptosis assessed by FACS in pLKO.1-shRNA control and CAL-51 NOXA knockdown stable cells treated with TAK-243 at the indicated concentration. (F) Western blot analysis showing dose response and time course of the effects of TAK-243 on UPR proteins in TNBC (CAL-51 and BT-549) and normal HCE-T cells, as assessed by immunoblotting for ATF4 and ATF6. Polyubiquitin (polyUb) indicates ubiquitin engagement in these cells. GAPDH was used as a loading control. For (E), error bars are S.E.M., n =3 and *p<0.05; ***p<0.001.

### TAK-243 induces unresolvable ER stress, mediated by ATF4-NOXA

Encouraged by the sensitivity to TAK-243 in the viability assays (Fig. 1B), we investigated whether inhibiting UBA1 was sufficient to induce cell death. Indeed, TAK-243 induced apoptotic cell death in TNBCs but had very little effect in normal tissue-derived HCE-T cells (Fig. 1C). BCL-2 family proteins mediate apoptosis (18), and examination of the level of these proteins demonstrated that TAK-243 did not downregulate key anti-apoptotic proteins in TNBC, nor did consistently upregulate the pro-apoptotic protein BIM. Interestingly, NOXA, another pro-apoptotic protein, was strongly induced in the TNBC models (12.5-fold in BT-549 and18.4-fold in CAL-51) but less so in HCET cells (2-fold) (Fig. 1D and Sup. Fig. S2).

To test whether NOXA was necessary for the observed effect of TAK-243, shRNA mediated depletion of NOXA was employed (Fig. 1E *left* panel). NOXA loss in TNBC cells led to mitigation of TAK-243-induced cell death (Fig. 1E, *right* panel and Sup. Fig. S2), demonstrating a critical role for NOXA in TAK-243 efficacy. NOXA is activated by both p53 and ER-stress, and disruption of protein homeostasis often induces ER-stress and the unfolded protein response (UPR), as well as the p53 response (19). To investigate the mechanism of NOXA upregulation, we evaluated both p53 response (Fig. 1D and Sup. Fig. S2) and ER stress response (Fig. 1F and Sup. Fig. S2). TAK-243 did not markedly induce p53 in neither the p53 mutant BT-549, MDA-MB-468 and HCC70 cells nor the p53 wild-type CAL-51 cells (Fig. 1D and Sup. Fig. S2). The UPR consists of three main pathway arms initiated by ER stress: 1) Protein kinase RNA-like reticulum kinase (PERK), which activates eukaryotic initiation factor 2 (eIF2 alpha) to repress general translation and promote specific translation of a set of mRNAs including activating transcription factor (ATF4); 2) Inositol-requiring enzyme 1 (IRE1), and 3) ATF6 (20). Interestingly, we found TAK-243 strongly induced ATF4 expression in every TNBC model analyzed, and to a lesser degree in variable models, ATF6 expression (Fig. 1F and Sup. Fig. S2).

To probe the potential functional role of ATF4 and ATF6 in NOXA upregulation, and response to TAK-243, ATF4 and ATF6 expression was reduced using specific shRNAs. While ATF6 knockdown did not impact NOXA induction or cell death, ATF4 depletion mitigated both NOXA upregulation and cell death following TAK-243 treatment (Fig. 2A). These data support a model where blocking UBA1 induces the UPR, leading to unresolvable ER-stress, ATF4-mediated NOXA upregulation, and apoptosis in TNBC (Fig. 2B).

**Figure 2.**
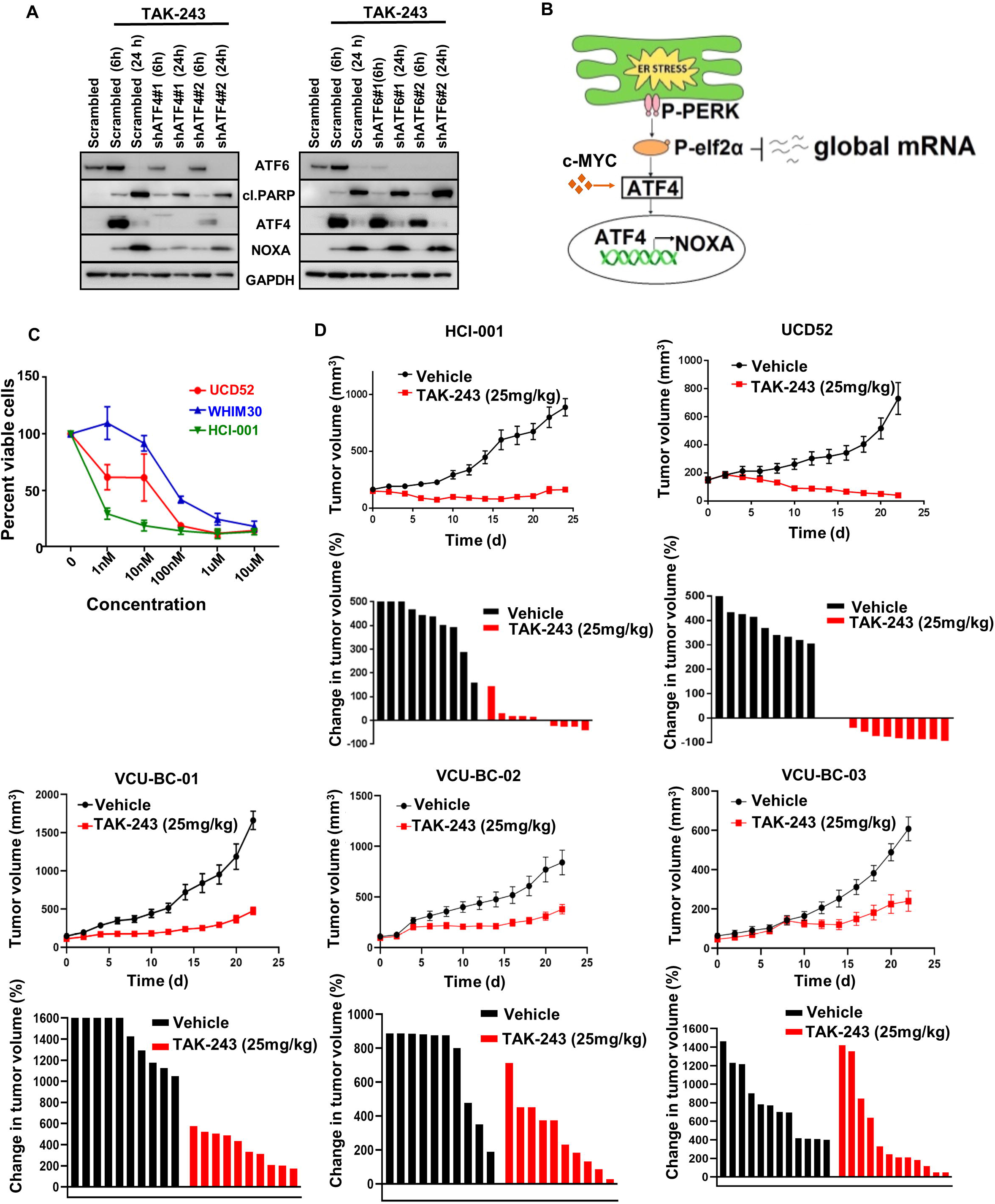
TAK-243 is highly effective in inhibiting TNBC tumor growth. (A) Western blot analysis demonstrating the role of ATF4 in the regulation of NOXA as assessed by immunoblotting. (B) Model for TAK-243 efficacy in TNBC. (C) Graph represents percent viable cells in *ex vivo* TNBC cell cultures (WHIM30, UCD52 and HCI-001) following 72 h treatment with TAK-243 at the indicated concentrations. (D) Anti-tumor activity of TAK-243 was assessed by tumor volume over time in mice bearing patient derived TNBC xenografts HCI-001, UCD52, VCU-BC-01, VCU-BC-02 and VCU-BC-03. Waterfall plot represents change in tumor volume percent of each tumor to their initial tumor size.

### TAK-243 induces some tumor regressions in TNBC patient-derived xenografts

We recently characterized several PDX models and corresponding cell lines (21); evaluation of TAK-243 in three PDX spheroid cell cultures demonstrated similar TAK-243 efficacy (Fig. 2C) as we found in the TNBC cell lines (Fig. 1B). Encouraged by the broad, albeit variable, sensitivity of TAK-243 across these TNBC PDX spheroid models, we treated female NSG mice bearing these TNBC PDX tumors and additional TNBC PDX models developed at VCU, with TAK-243 (16). (Sup. Table 3 and 4). Treatment was sufficient to induce strong responses with overt tumor regression in one PDX model and with strong, albeit a more mixed response, in the other four PDX models (Fig. 2D). Immunoblot analysis of control and TAK-243 treated tumor tissues further corroborated our *in vitro* results (Sup. Fig. S3). Immunohistochemical analysis of the expression of the cell death marker cl. Caspase 3 (CC3) further validated our findings of increased cell death following TAK-243 treatment compared to vehicle-treated PDX mice (Sup. Fig. S3). Treatment tolerability was indicated by weight maintenance throughout the study (Sup. Fig. S4). Overall, these data are consistent with the *in vitro* findings from the TNBC cell lines, demonstrating a high degree of activity across TNBC models through targeting of UBA1, which can be recapitulated with the small molecule UBA1 inhibitor, TAK-243.

### TAK-243 decreases primary metastases in a metastatic mouse model of TNBC

Metastasis is responsible for almost all TNBC deaths. TNBC metastasizes to the lymph nodes, liver, lung, bone, and brain. We evaluated the effect of TAK-243 on metastatic growth using an intracardiac MDA-MB-231-tomato-luciferase injection model. Drug treatment did not begin until 10 days after injection of the cells allowing time for the cells to extravasate to all organs and begin to grow. Importantly, we observed an overall decrease in the metastatic burden of the NSG mice treated with TAK-243 [(25mg/kg), Fig 3A & B)] in primary metastatic regions such as the lung, liver, and bones as well as the ovaries, and kidney (Fig 3C & Sup. Fig S5). These data demonstrate that TAK-243 decreases TNBC metastatic burden.

**Figure 3.**
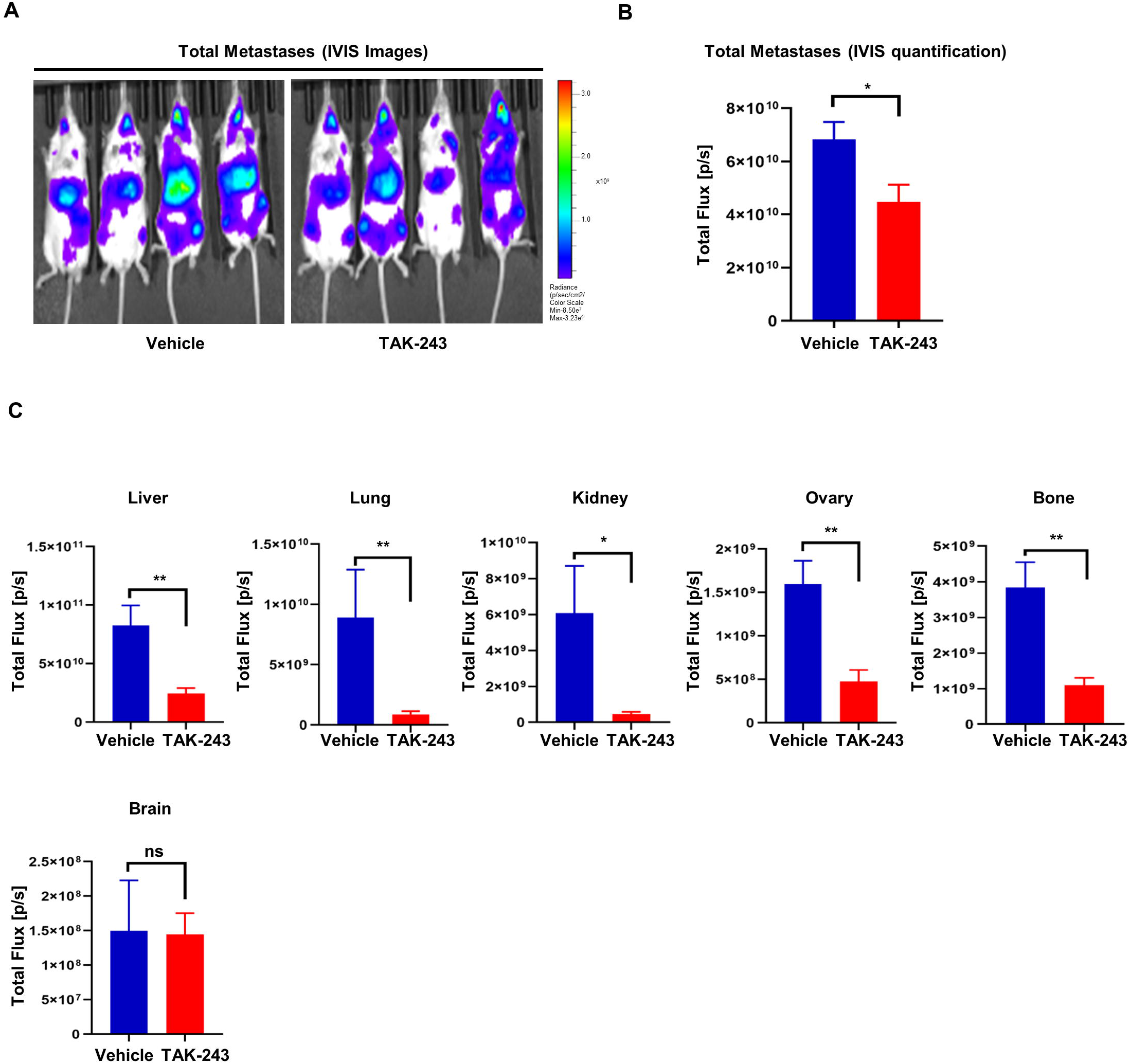
TAK-243 decreases metastases in TNBC. (A) Total metastases were imaged *in vivo* in MDA-MB-231 tomato-luciferase model. (B) Graph represents total metastases quantified *in vivo* in MDA-MB-231 tomato-luciferase model. (C) Graph represents metastases quantified *ex vivo* in the organs (liver, lung, kidney, ovaries, bone, and brain) of vehicle and TAK-243 treated mice (*p<0.05; **p<0.01, ns: not significant).

### c-MYC is a functional biomarker for TAK-243 activity across TNBC models

As high c-MYC levels are detected often in TNBC (22) and we recently demonstrated that n-MYC, a closely related paralog of c-MYC, sensitizes cancer cells to NOXA-mediated toxicity (23), we asked whether c-MYC expression could be a potential factor in TAK-243 efficacy in TNBC. To do this, we analyzed c-MYC expression across our panel of TNBC cell lines and normal-tissue derived cells. We found an increase in c-MYC expression between the sensitive TNBCs and the normal-tissue derived cells (Fig 4A; *left* and *right* panel). Moreover, we detected increased expression of c-MYC in the TAK-243 sensitive TNBCs compared to insensitive TNBCs (Fig 4A; *left* and *right* panel). We also noted that in the PDXs, the best responders to TAK-243 (HCI-001 and UCD52) had higher levels of c-MYC compared to the other PDXs tested (VCU-BC-01, VCU-BC-02 and VCU-BC-03) (Fig 4B), suggesting that c-MYC expression might correlate with sensitivity to TAK-243 in *in vivo* as well.

**Figure 4.**
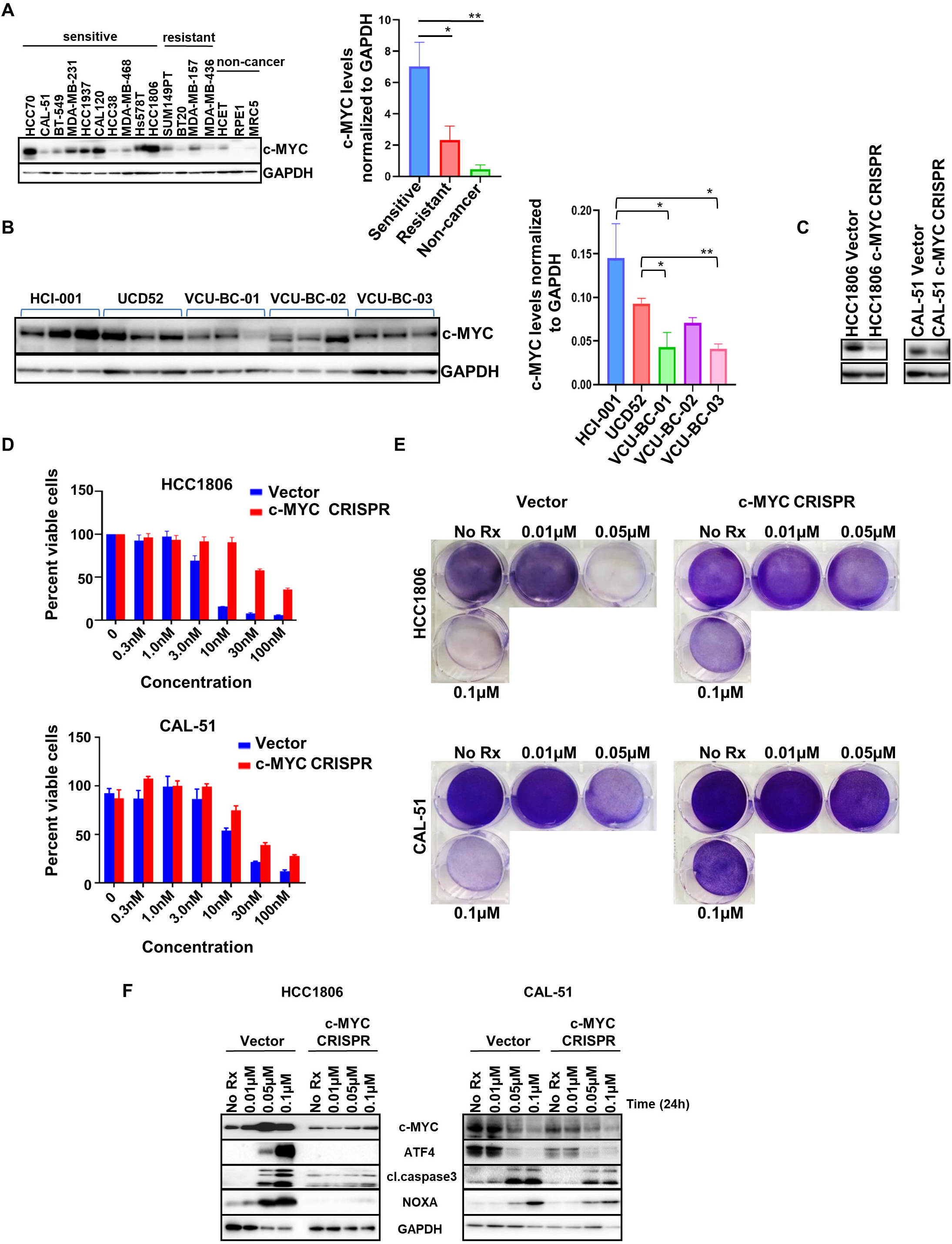
c-MYC mediates sensitivity to TAK-243 in TNBC. (A) Western blot analysis showing basal levels of c-MYC in TNBC-sensitive to TAK-243, resistant to TAK-243 and normal cell lines (*left* panel). Right panel indicates c-MYC levels normalized to GAPDH in the sensitive, resistant and non-cancer cell lines (*p<0.05; **p<0.01). (B) Immunoblot showing c-MYC expression in PDX tumor lysates. Right panel indicates c-MYC levels normalized to GAPDH in the PDX tumor lysates (*p<0.05; **p<0.01). (C) Immunoblot showing c-MYC expression in plenti CRISPR v2 (control) and plenti CRISPR c-MYC stable lines. (D) Graph represents percent viable cells assessed by Cell Titer-Glo in two plenti CRISPR v2 (control, blue) and plenti CRISPR c-MYC stable lines (red) following 72 h treatment with TAK-243 at the indicated concentrations. (E) Crystal violet assay in plenti CRISPR v2 (vector) and plenti CRISPR c-MYC stable lines treated for 4d with TAK-243 at the indicated concentrations. (F) Immunoblot showing expression of cl.PARP, CC3 and NOXA in plenti CRISPR v2 (vector) and plenti CRISPR c-MYC stable cells treated with TAK-243 for 24h at the indicated concentrations.

To determine if c-MYC can modulate TAK-243 sensitivity, we knocked out c-MYC in HCC1806 and CAL-51 cells using CRISPR/Cas9 technology (Fig 4C). Indeed, downregulation of c-MYC expression decreased the sensitivity to TAK-243 (Fig. 4D and E). Consistent with the ability of MYC to induce an ER-stress response (24) and ATF4 expression (25-27), cells with reduced c-MYC expression had decreased expression of ATF4 and NOXA following TAK-243 treatment. Consistent with these findings, there was a reduction in TAK-243 mediated cell death in the knock out cells, as determined by CC3 expression (Fig 4F). To confirm the involvement of c-MYC in TAK-243 sensitivity in TNBC, we overexpressed c-MYC with lentiviral producing c-MYC-expressing plasmids. Consistent with the CRISPR experiment results, exogenous c-MYC expression in CAL-51 lines increased the sensitivity to TAK-243 even further (Fig. 5A-B). Western blot analyses revealed that c-MYC increased levels of ATF4 and NOXA (Fig. 5C); consistent with this observation, cell death markers were increased in the c-MYC overexpressing cells following TAK-243 (Fig. 5C). Altogether, these data suggest that c-MYC by activating the ATF4/NOXA axis, can cooperate with TAK-243 in inducing stress response and cell death, and therefore c-MYC may serve as a functional biomarker for response to TAK-243 in TNBC.

**Figure 5.**
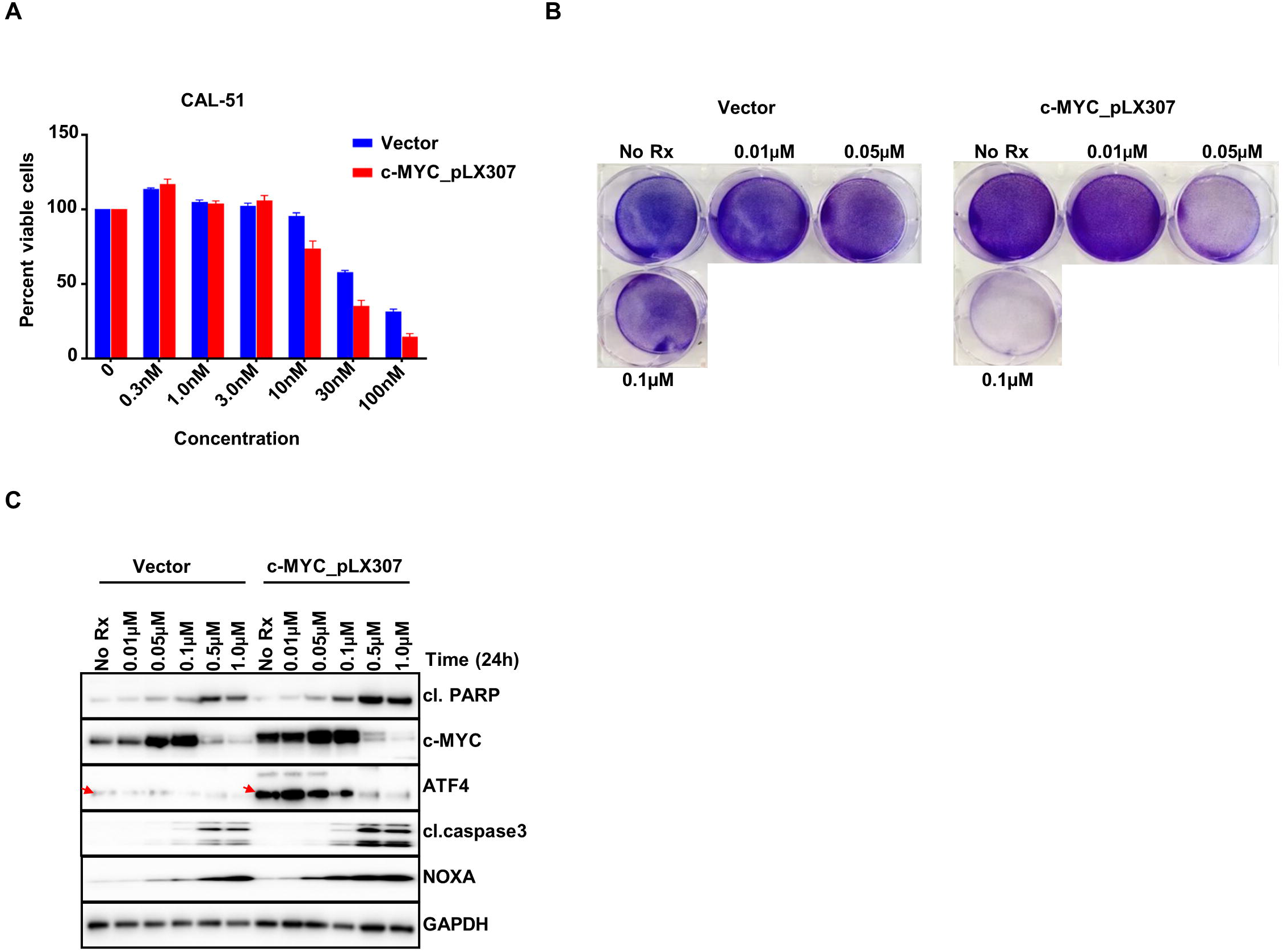
c-MYC overexpression increases sensitivity to TAK-243 in TNBC. (A) Graph represents percent viable cells assessed by Cell Titer-Glo in lentiviral pLX307 (vector) and c-MYC_pLX307 expressing CAL-51 cells following 72 h treatment with TAK-243 at the indicated concentrations. (B) Crystal violet assay in lentiviral pLX307 (vector) and c-MYC_pLX307 expressing CAL-51 cells treated for 3d with TAK-243 at the indicated concentrations. (C) Immunoblot showing expression of cl.PARP, CC3 and NOXA in lentiviral pLX307 (vector) and c-MYC_pLX307 expressing CAL-51 treated with TAK-243 for 24h at the indicated concentrations.

## Discussion

TNBC continues to be routinely treated with chemotherapy where responses are variable; as a result, TNBC remains the most fatal breast cancer subtype, despite occurring in younger women.

Full genome CRISPR/Cas9 screening offers tangible advantages to past screening efforts by revealing a greater number of phenotypes through improved penetrance, which is a characteristic of genome editing itself compared to, for instance, screening with short-interfering (si) RNA or short-hairpin (sh) RNA libraries (13). We hypothesized that this technique could reveal a drug target that we could capitalize on, given the growing number of clinically available targeted therapies emerging that are designed to interfere with genes outside the traditional kinome.

Cellular protein homeostasis is maintained through a careful balance of protein synthesis and degradation. The consequence of disruptions in this homeostasis causes diverse human diseases, including cancer (16, 17). Protein synthesis is mainly countered in mammalian cells by the UPS where the protein ubiquitin “tags” proteins for degradation in several cellular proteasomes (17). Here, we found that blocking UBA1, the enzyme responsible for adenylating and capturing a ubiquitin molecule to initiate ubiquitination of cellular proteins (28), was highly toxic to various TNBC models and led to ER-stress and NOXA upregulation, initiating apoptosis.

While proteasome inhibitors have long been investigated as anti-cancer agents (29), UBA1 inhibitors have taken longer to be developed. Schimmer and colleagues (28) demonstrated that UBA1 was a *bona fide* drug target in several hematological cancers several years ago. This hypothesis became testable in hematological cancers with the recent discovery and characterization of TAK-243, a specific and potent UBA1 inhibitor (16). Indeed, the same group found that acute myeloid leukemia (AML) models are significantly sensitivity to TAK-243 (15, 30).

Here, we demonstrate that TAK-243 has anti-tumor activity also in solid tumors. Indeed, we found that most TNBCs were sensitive to TAK-243, while normal-tissue derived cells were relatively refractory to TAK-243, being roughly an order of magnitude less sensitive than the TNBC cell lines. TAK-243 was sufficient to uniformly block tumor growth in different TNBC PDXs including the ones developed at VCU. (Fig. 2D). Additionally, TAK-243 decreased metastatic burden of MDA-MB-231-tomato-luc cells in different organs (Fig 4) demonstrating TAK-243 can decrease both the primary tumor and tumors at various metastatic sites.

Moreover, we uncovered the oncogenic transcription factor, c-MYC, as both correlating with TAK-243 sensitivity and having a causative role in TAK-243 sensitivity. c-MYC is an important oncogenic transcription factor in several cancers including TNBC (31, 32). In TNBC, c-MYC contributes to chemotherapy resistance (33). As such, c-MYC is a premiere drug target in TNBCs and other cancers but has been historically difficult to target directly due to its protein structure and other characteristics (34). One strategy is to target processes or druggable proteins that are preferentially active in c-MYC expressing TNBCs. For instance, PIM1 is higher in c-MYC-expressing TNBCs (35) as is fatty acid oxidation (36) and targeting both these processes in these cancers are effective in preclinical models. While PIM inhibitors have been tested clinically (e.g. NCT01588548), the lack of objective responses across hematological and solid tumors have hindered development (37).

In addition to its roles in supporting proliferation and growth, c-MYC increases ER stress (24, 38) and in particular has an intimate relationship with ATF4 (25). Indeed, both c-MYC and n-MYC have been demonstrated to upregulate ATF4, which may involve the kinase GNC2 (25). Our finding that the expression of c-MYC was important in determining TAK-243 sensitivity is consistent with a recent TAK-243 study in the hematological cancer DLBCL (39), and conceptually with our recent report with n-MYC (23), where we demonstrated n-MYC sensitizes to NOXA-mediated death in neuroblastoma. We propose here that TAK-243 treatment can capitalize on the MYC effects on ER-stress, by inducing unresolved ER stress preferentially in c-MYC expressing TNBCs, with cells dying through a NOXA-dependent mechanism. While there is an ER-stress response in the sensitive TNBCs without c-MYC expression, it is not as substantial and therefore there is a parallel reduction in cell death and efficacy of TAK-243. Lastly, we found in the normal tissue-derived cells a relatively tempered ER-stress response, which corresponded with less toxicity. Thus, c-MYC dependent ATF4/NOXA induction increases sensitivity of TNBC models to TAK-243 and c-MYC may act as a biomarker for response. Of note, a diagnostic c-MYC immunohistochemical staining has been developed for hematological cancers (40), and this could potentially support the use of c-MYC in future clinical studies of TAK-243 or newer UBA1 inhibitors in TNBC.

Interestingly, and consistent with our findings, other studies support the role of the UPS in TNBC tumor growth: Lieberman and colleagues previously noted the efficacy of proteasome inhibitors in mouse models of TNBC (12). Furthermore, UBR5, the E3 ubiquitin ligase, is overexpressed in TNBC and genetic targeting of UBR5 results in tumor growth inhibition (41). UBR5 is currently not druggable and UBA1 inhibitors may be preferential to proteasome inhibitors for several reasons. First, proteasome inhibitors can be quite toxic to some normal cells (42-44), and while UBA1 inhibitors may also demonstrate normal cell toxicity, our data suggest a therapeutic window. Secondly, Kisselev and colleagues demonstrated targeting additional proteasome sites over what the clinically advanced proteasome inhibitors achieve is necessary for efficacy (45). Thirdly, there is transcriptional feedback to maintain proteasome activity, which can be mediated by NRF1 (46). Thus, it appears that TNBCs are vulnerable to targeting of the UPS and targeting the UPS at the UBA1 stage may offer a better chance of a durable target and a therapeutic window.

In summary, we have demonstrated that targeting UBA1 with TAK-243 in high c-MYC-expressing TNBCs has preclinical activity and may provide an effective alternative therapeutic approach over chemotherapy. Thus, TAK-243 may be a valuable targeted therapy approach across diverse TNBC models.

## Material and Methods

### CRISPR Screening

TNBC cell lines BT-549 and MDA-MB-468 were infected with a Lentiviral genome-wide CRISPR-guide RNA library of unique sgRNAs targeting 20,000 genes (4 sgRNAs per gene). Cells were selected in puromycin and cultured for 21 days before isolating genomic DNA and quantifying sgRNA in surviving cells by deep sequencing. This full genomic screen was followed by screening nine TNBCs-cell lines (MDA-MB-468, MDA-MB-231, Hs578T, SUM149PT, BT-549, HCC1806, HCC70, MDA-MB-436, HCC1937) with 10 sgRNAs against UBA1.

### Cell Lines

The TNBC cell lines in this study were cultured in their respective media with 10% FBS in the presence of 1 μg/mL penicillin and streptomycin. BT-549, HCC1937, HCC70, CAL-51, HCC1806, BT20, MDA-MB-157, MDA-MB-436 and RPE1 were grown in RPMI while MDA-MB-231, MDA-MB-468, HCC38, SUM149PT and HCE-T were cultured in DMEM/F12. DMEM was used to culture CAL-120 and MRC5.

### Reagents and Antibodies

TAK-243 (MLN 7243) was purchased from Chemietek (CT-M7243) for *in vitro* and *in vivo* studies. The antibodies used in this study were purchased from Cell Signaling and Santa Cruz. All primary antibodies were used at 1:1000 dilutions. Corresponding HRP-conjugated secondary antibodies were used at 1:5000 dilution.

### Cell Viability Assay

For Cell Titer-Glo experiments, 2000-3000 cells were seeded in 96-well flat-bottom black plates and treated with increasing concentration of drug for 72h, as previously described (47). Following drug treatment, 25 μL of CellTiter-Glo (Promega), was added and read on a Centro LB 960 microplate luminometer (Berthold Technologies). For crystal violet assay, cells were seeded at 50,000 cells/well of 6-well plate. Cells were treated with increasing concentration of drug as indicated and incubated until no treatment wells were confluent. Cells were then fixed with 50% glutaraldehyde and stained with 0.1% crystal violet (Sigma-Aldrich) and visualized.

### FACS Apoptosis Assay

3×10^5^ cells seeded per well in six well plates were drugged with the desired concentration of TAK-243 for 24h, or left untreated, as previously described (48). Cells were incubated with propidium iodide and annexin V–Cy5 (BD Biosciences) together for 15 min and assayed on a Guava easyCyte flow cytometer (Millipore Sigma). Analysis was performed using FlowJo. Cells stained positive for annexin V-Cy5 and annexin V + propidium were counted as apoptotic.

### Western Blot Analysis

Cell lysates prepared in NP-40 lysis buffer (20 mM Tris, 150 mM NaCl, 1% Nonidet P-40, 1 mM EDTA, 1 mM EGTA, 10% glycerol, and protease and phosphatase inhibitors), were incubated on ice for 30 min before centrifugation at high speed for 10 min at 4°C. Tumor lysates were homogenized with Tissuemiser (Fisher Scientific) in lysis buffer. Equal amounts of detergent-soluble lysates were resolved using the NuPAGE Novex Midi Gel system on 4–12% Bis–Tris gels (Invitrogen). Proteins were transferred to PVDF membranes (PerkinElmer) and blocked in 5% nonfat milk in PBS. Blots were probed with the primary antibodies overnight and later with species appropriate HRP conjugated secondary antibodies. Chemiluminescence was detected with the Syngene G:Box camera (Synoptics).

### Immunohistochemical (IHC) Staining

IHC staining for CC3 (1:500, Cell Signaling #9664) was performed in the VCU Tissue and Data Acquisition and Analysis Core with the Leica Bond RX autostainer using heat-induced epitope retrieval buffer 2 (Leica, EDTA pH 8.0) for 20 min with antibody incubation for 45min. Stained slides were then imaged on the Vectra Polaris (PerkinElmer) and scored. H-score was determined by multiplication of the percentage of cells with staining intensity ordinal value (3 x % of cells with 3+ intensity) + (2 x % of cells with 2+ intensity) + (1 x % of cells with 1+ intensity) = H-Score.

### Vector Construction and Stable Cell Lines

For the short-hairpin RNA (shRNA) experiments, shATF4 and shATF6 purchased from Dharmacon and the shNOXA previously described (48) was used. shRNA designed against a scramble sequence (MISSION pLKO.1-shRNA control plasmid DNA) served as the control. For the c-MYC over expression, MYC_pLX307 plasmid (Addgene#98363) and pLX307 empty vector (Addgene#117734) were used. Cells were transduced with plasmid containing viral particles that were generated in 293T cells and collected over 48h, as previously described (47). The pLKO.1 puromycin resistant vector backbone served as the basis for cell selection in puromycin following infection. plenti CRISPR v2 virus against sgRNA targeting c-MYC (Genscript) and plenti CRISPR v2 virus (control) was commercially made to order from Science Exchange. Cells were then infected with viral particles and selected in puromycin to generate c-MYC KD stable lines.

### PDX and *ex vivo* Studies

All animal experiments were conducted in accordance with a protocol approved by VCU Institutional Animal Care and Use Committee. PDX models HCI-001, WHIM30 and UCD52 were obtained from the University of Utah/Huntsman Cancer Institute, Washington University, St. Louis, and the University of Colorado, respectively, and expanded in NSG mice obtained and bred by VCU Cancer Mouse Models Core (CMMC). The VCU-BC PDX (Sup Table 3) were developed by VCU CMMC with TNBC tumor samples obtained from VCU Tissue Data Acquisition and Analysis Core (collected under a protocol approved by VCU Institutional Review Board #HM2471). For *ex vivo* studies, PDX tumors were removed, finely chopped, and digested in DMEM/F12 containing 5% FBS, 300U/ml collagenase (Sigma) and 100 U/ml hyaluronidase (Sigma). Tissue was then suspended in ((NH4)Cl) followed by trypsinization to generate suspensions of single cells, which were previously transduced with lentivirus (BLIV101PA-1, Systems Biosciences) encoding GFP and Luciferase. Cell suspensions were plated at 25,000 cells/well in 96-well plates in M87 medium, followed by treatment with varying concentrations of TAK-243 in triplicate. After 72h of treatment, luciferin was added to each well (10% of total volume per well) and plates were imaged using the IVIS Spectrum In Vivo Imaging System (PerkinElmer). Cell viability was measured as total photon flux per second, and drug response was evaluated based on percent of vehicle viability (49). For *in vivo* studies, 500,000 cells, suspended 1:1 in Matrigel, were injected into the fourth mammary fat pads of experimental female NSG mice. Mice with tumor volumes of ∼150– 200 mm3 were randomized into two groups: TAK-243 treatment and control treatment (vehicle only) and dosed intravenously. TAK-243 was formulated in the vehicle consisting of: 25 mM HCl, 20% 2-hydroxy propyl-β-cyclodextrin (Sigma-Aldrich). Animals were treated with 25mg/kg of TAK-243 biweekly (Mon & Thur) for 3 wk. Tumors were measured daily by caliper, in two dimensions (length and width), and tumor volume was calculated with the formula v = l × (w)2 (π/6), where v is the tumor volume, l is the length, and w is the width (the smaller of the two measurements). Following the final treatment, the tumors were harvested after 2h and snap frozen in liquid nitrogen for pharmacodynamic studies.

### Experimental Metastasis Study

MDA-MB-231-tomato-luciferase cells (1×10^5^ for endpoint study) in 100-µL sterile PBS were injected into the left cardiac ventricle of female 5-week-old NSG mice as described previously (50) and *in vivo* imaging was performed (IVIS Spectrum, PerkinElmer) immediately to verify widespread seeding of tumor cells. Mice were randomized using Multi-Task program on day 10 into two groups: TAK-243 treatment and controls and dosed intravenously with TAK-243 formulated in 25 mM HCl, 20% 2-hydroxy propyl-β-cyclodextrin (Sigma-Aldrich) or vehicle alone. Animals were treated with 25mg/kg of TAK-243 biweekly (Mon & Thur) for 2 wks. Bioluminescence (radiance/sec) emitted from the cells were quantified using the IVIS Spectrum and Living Image software (PerkinElmer) of the animals. For the endpoint study, mice were euthanized, and bioluminescence of harvested organs (kidneys, lung, ovaries, liver, brain, and skeleton) were quantified by ex vivo imaging and analysis.

### Statistical Analyses

Unpaired student’s t-test (two-tailed) was performed for figures 1E & 3C; Mann Whitney and Welch’s unpaired t test was performed for figures 4A & 4B using GraphPad Prism. Differences considered to be significant if p< 0.05.

## Acknowledgments

Services and products in support of the research project were generated by the Virginia Commonwealth University Cancer Mouse Models Core and Tissue and Data Acquisition and Analysis Core Laboratory, supported, in part, with funding to the Massey Cancer Center from NIH-NCI Cancer Center Support Grant P30 CA016059. This work was supported by a VCU Massey Pilot grant awarded to A.C.F., J.E.K. and M.G.D.

## Acknowledgements

Services and products in support of the research project were generated by the Virginia Commonwealth University Cancer Mouse Models Core Laboratory and Tissue Data Acquisition and Analysis Core, supported, in part, with funding to the Massey Cancer Center from NIH-NCI Cancer Center Support Grant P30 CA016059. This work was supported by a Massey Cancer Center Pilot Grant (A.C.F., S.B. and M.G.D.).

## Study Approval

All mouse experiments were approved and performed in accordance with the Institutional Animal Care and Use Committee at VCU (IACUC protocol AD10001048 and AD10001247).

## Supplementary table legends

**Table 1**. Top 400 genes (approximately 2% of total genes targeted) by STARS scores.

**Table 2**. Ubiquitination related hits and their scores in the primary screen analysis.

**Table 3**. Details of VCU PDX models.

## Supplementary figures

**Figure 1. UBA1 expression levels in TNBC lines**. Graph represents average Log Fold Change (LFC) in the expression of guide RNAs against UBA1 in nine TNBC cell lines. Average LFC was calculated from LFC of 10 clones/sgRNA of UBA1 in the CRISPR analysis.

**Figure 2. TAK-243 induced cell death in TNBC**. (A) Western blot analysis showing dose response and time course of the effects of TAK-243 on ubiquitination and apoptosis in TNBC (MDA-MB-468 and HCC70) and normal HCE-T cells, as assessed by immunoblotting for p53, MCL-1, Bcl-xL, BIM, PUMA, cleaved Caspase 3 (CC3) and NOXA. GAPDH was used as a loading control. (B) Western blot analysis showing dose response and time course of the effects of TAK-243 on UPR proteins in TNBC (MDA-MB-468 and HCC70) and normal HCE-T cells, as assessed by immunoblotting for ATF4 and ATF6. Polyubiquitin (polyUb) indicates ubiquitin engagement in these cells. GAPDH was used as a loading control. (C) Polyubiquitin (polyUb) indicates ubiquitin engagement in these cells. GAPDH was used as a loading control. (C) FACS analysis demonstrating Annexin-V-Cy5 and Propidium Iodide staining in pLKO.1-shRNA control and CAL-51 NOXA knockdown stable cells following 24h treatment with TAK-243 at the indicated concentrations.

**Figure 3. TAK-243 induces cell death *in vivo***. (A) Western blot analysis showing effects of TAK-243 on UPR and apoptotic proteins in PDX HCI-001 mice treated with TAK-243, as assessed by immunoblotting for polyubiquitin (polyUb), ATF4, ATF6, CC3 and NOXA. GAPDH was used as a loading control. (B) Representative images of cl. caspase 3 (CC3) staining of vehicle (DMSO) or TAK-243 (25mg/kg, biweekly for 3 weeks) treated PDX tumors (UCD52). Total magnification 20x (scale bar: 50µm). Graph indicates H-Score analysis of the staining in the vehicle and TAK-243 treated tumor tissues (**p<0.01).

**Figure 4. TAK-243 has minimal effect on animal weight**. Graph displays the mouse weights of NSG mice bearing (A) PDX HCI-001, (B) PDX UCD52, (C) VCU-BC-01 (D) VCU-BC-02 and (E) VCU-BC-03 throughout the efficacy studies.

**Figure 5. TAK-243 decreases metastases in primary organs**. Images represent total metastases quantified *ex vivo* in the liver and lung of the vehicle and TAK-243 treated mice.

## References

1. Foulkes WD, Smith IE, Reis-Filho JS. Triple-negative breast cancer. N Engl J Med. 2010;363(20):1938–48.

2. Jhan JR, Andrechek ER. Triple-negative breast cancer and the potential for targeted therapy. Pharmacogenomics. 2017;18(17):1595–609.

3. Rottenberg S, Jaspers JE, Kersbergen A, van der Burg E, Nygren AO, Zander SA, et al. High sensitivity of BRCA1-deficient mammary tumors to the PARP inhibitor AZD2281 alone and in combination with platinum drugs. Proc Natl Acad Sci U S A. 2008;105(44):17079–84.

4. Fong PC, Boss DS, Yap TA, Tutt A, Wu P, Mergui-Roelvink M, et al. Inhibition of poly(ADP-ribose) polymerase in tumors from BRCA mutation carriers. N Engl J Med. 2009;361(2):123–34.

5. Loibl S, O’Shaughnessy J, Untch M, Sikov WM, Rugo HS, McKee MD, et al. Addition of the PARP inhibitor veliparib plus carboplatin or carboplatin alone to standard neoadjuvant chemotherapy in triple-negative breast cancer (BrighTNess): a randomised, phase 3 trial. The lancet oncology. 2018;19(4):497–509.

6. Clamp A, Jayson G. PARP inhibitors in BRCA mutation-associated ovarian cancer. Lancet Oncol. 2015;16(1):10–2.

7. Millis SZ, Gatalica Z, Winkler J, Vranic S, Kimbrough J, Reddy S, et al. Predictive Biomarker Profiling of > 6000 Breast Cancer Patients Shows Heterogeneity in TNBC, With Treatment Implications. Clin Breast Cancer. 2015;15(6):473–81 e3.

8. Castroviejo-Bermejo M, Cruz C, Llop-Guevara A, Gutierrez-Enriquez S, Ducy M, Ibrahim YH, et al. A RAD51 assay feasible in routine tumor samples calls PARP inhibitor response beyond BRCA mutation. EMBO molecular medicine. 2018;10(12).

9. Kwa MJ, Adams S. Checkpoint inhibitors in triple-negative breast cancer (TNBC): Where to go from here. Cancer. 2018;124(10):2086–103.

10. Bassey-Archibong BI, Hercules SM, Rayner LGA, Skeete DHA, Smith Connell SP, Brain I, et al. Kaiso is highly expressed in TNBC tissues of women of African ancestry compared to Caucasian women. Cancer causes & control : CCC. 2017;28(11):1295–304.

11. Yau EH, Kummetha IR, Lichinchi G, Tang R, Zhang Y, Rana TM. Genome-Wide CRISPR Screen for Essential Cell Growth Mediators in Mutant KRAS Colorectal Cancers. Cancer Res. 2017;77(22):6330–9.

12. Petrocca F, Altschuler G, Tan SM, Mendillo ML, Yan H, Jerry DJ, et al. A genome-wide siRNA screen identifies proteasome addiction as a vulnerability of basal-like triple-negative breast cancer cells. Cancer Cell. 2013;24(2):182–96.

13. Braso-Maristany F, Filosto S, Catchpole S, Marlow R, Quist J, Francesch-Domenech E, et al. PIM1 kinase regulates cell death, tumor growth and chemotherapy response in triple-negative breast cancer. Nat Med. 2016;22(11):1303–13.

14. Groen EJN, Gillingwater TH. UBA1: At the Crossroads of Ubiquitin Homeostasis and Neurodegeneration. Trends Mol Med. 2015;21(10):622–32.

15. Barghout SH, Patel PS, Wang X, Xu GW, Kavanagh S, Halgas O, et al. Preclinical evaluation of the selective small-molecule UBA1 inhibitor, TAK-243, in acute myeloid leukemia. Leukemia. 2018.

16. Hyer ML, Milhollen MA, Ciavarri J, Fleming P, Traore T, Sappal D, et al. A small-molecule inhibitor of the ubiquitin activating enzyme for cancer treatment. Nat Med. 2018;24(2):186–93.

17. Xu W, Lukkarila JL, da Silva SR, Paiva SL, Gunning PT, Schimmer AD. Targeting the ubiquitin E1 as a novel anti-cancer strategy. Curr Pharm Des. 2013;19(18):3201–9.

18. Hata AN, Engelman JA, Faber AC. The BCL2 Family: Key Mediators of the Apoptotic Response to Targeted Anticancer Therapeutics. Cancer Discov. 2015;5(5):475–87.

19. Lee G, Oh TI, Um KB, Yoon H, Son J, Kim BM, et al. Small-molecule inhibitors of USP7 induce apoptosis through oxidative and endoplasmic reticulum stress in cancer cells. Biochem Biophys Res Commun. 2016;470(1):181–6.

20. Szegezdi E, Logue SE, Gorman AM, Samali A. Mediators of endoplasmic reticulum stress-induced apoptosis. EMBO Rep. 2006;7(9):880–5.

21. Alzubi MA, Turner TH, Olex AL, Sohal SS, Tobin NP, Recio SG, et al. Separation of breast cancer and organ microenvironment transcriptomes in metastases. Breast Cancer Res. 2019;21(1):36.

22. Klauber-DeMore N, Schulte BA, Wang GY. Targeting MYC for triple-negative breast cancer treatment. Oncoscience. 2018;5(5-6):120–1.

23. Ham J, Costa C, Sano R, Lochmann TL, Sennott EM, Patel NU, et al. Exploitation of the Apoptosis-Primed State of MYCN-Amplified Neuroblastoma to Develop a Potent and Specific Targeted Therapy Combination. Cancer Cell. 2016;29(2):159–72.

24. Zhang T, Li N, Sun C, Jin Y, Sheng X. MYC and the unfolded protein response in cancer: synthetic lethal partners in crime? EMBO Mol Med. 2020;12(5):e11845.

25. Tameire F, Verginadis, II, Leli NM, Polte C, Conn CS, Ojha R, et al. ATF4 couples MYC-dependent translational activity to bioenergetic demands during tumour progression. Nat Cell Biol. 2019;21(7):889–99.

26. Qing G, Li B, Vu A, Skuli N, Walton ZE, Liu X, et al. ATF4 regulates MYC-mediated neuroblastoma cell death upon glutamine deprivation. Cancer Cell. 2012;22(5):631–44.

27. Xia Y, Ye B, Ding J, Yu Y, Alptekin A, Thangaraju M, et al. Metabolic Reprogramming by MYCN Confers Dependence on the Serine-Glycine-One-Carbon Biosynthetic Pathway. Cancer Res. 2019;79(15):3837–50.

28. Xu GW, Ali M, Wood TE, Wong D, Maclean N, Wang X, et al. The ubiquitin-activating enzyme E1 as a therapeutic target for the treatment of leukemia and multiple myeloma. Blood. 2010;115(11):2251–9.

29. Gandolfi S, Laubach JP, Hideshima T, Chauhan D, Anderson KC, Richardson PG. The proteasome and proteasome inhibitors in multiple myeloma. Cancer Metastasis Rev. 2017;36(4):561–84.

30. Barghout SH, Schimmer AD. The ubiquitin-activating enzyme, UBA1, as a novel therapeutic target for AML. Oncotarget. 2018;9(76):34198–9.

31. Xu J, Chen Y, Olopade OI. MYC and Breast Cancer. Genes Cancer. 2010;1(6):629–40.

32. Felsher DW, Bishop JM. Reversible tumorigenesis by MYC in hematopoietic lineages. Mol Cell. 1999;4(2):199–207.

33. Lee KM, Giltnane JM, Balko JM, Schwarz LJ, Guerrero-Zotano AL, Hutchinson KE, et al. MYC and MCL1 Cooperatively Promote Chemotherapy-Resistant Breast Cancer Stem Cells via Regulation of Mitochondrial Oxidative Phosphorylation. Cell Metab. 2017;26(4):633–47 e7.

34. Allen-Petersen BL, Sears RC. Mission Possible: Advances in MYC Therapeutic Targeting in Cancer. BioDrugs. 2019;33(5):539–53.

35. Horiuchi D, Camarda R, Zhou AY, Yau C, Momcilovic O, Balakrishnan S, et al. PIM1 kinase inhibition as a targeted therapy against triple-negative breast tumors with elevated MYC expression. Nat Med. 2016;22(11):1321–9.

36. Camarda R, Zhou AY, Kohnz RA, Balakrishnan S, Mahieu C, Anderton B, et al. Inhibition of fatty acid oxidation as a therapy for MYC-overexpressing triple-negative breast cancer. Nat Med. 2016;22(4):427–32.

37. Cortes J, Tamura K, DeAngelo DJ, de Bono J, Lorente D, Minden M, et al. Phase I studies of AZD1208, a proviral integration Moloney virus kinase inhibitor in solid and haematological cancers. Br J Cancer. 2018;118(11):1425–33.

38. Hart LS, Cunningham JT, Datta T, Dey S, Tameire F, Lehman SL, et al. ER stress-mediated autophagy promotes Myc-dependent transformation and tumor growth. J Clin Invest. 2012;122(12):4621–34.

39. Best S, Hashiguchi T, Kittai A, Bruss N, Paiva C, Okada C, et al. Targeting ubiquitin-activating enzyme induces ER stress-mediated apoptosis in B-cell lymphoma cells. Blood Adv. 2019;3(1):51–62.

40. Kluk MJ, Ho C, Yu H, Chen BJ, Neuberg DS, Dal Cin P, et al. MYC Immunohistochemistry to Identify MYC-Driven B-Cell Lymphomas in Clinical Practice. Am J Clin Pathol. 2016;145(2):166–79.

41. Liao L, Song M, Li X, Tang L, Zhang T, Zhang L, et al. E3 Ubiquitin Ligase UBR5 Drives the Growth and Metastasis of Triple-Negative Breast Cancer. Cancer Res. 2017;77(8):2090–101.

42. Kharel P, Uprety D, Chandra AB, Hu Y, Belur AA, Dhakal A. Bortezomib-Induced Pulmonary Toxicity: A Case Report and Review of Literature. Case Rep Med. 2018;2018:2913124.

43. Boyer JE, Batra RB, Ascensao JL, Schechter GP. Severe pulmonary complication after bortezomib treatment for multiple myeloma. Blood. 2006;108(3):1113.

44. Argyriou AA, Iconomou G, Kalofonos HP. Bortezomib-induced peripheral neuropathy in multiple myeloma: a comprehensive review of the literature. Blood. 2008;112(5):1593–9.

45. Weyburne ES, Wilkins OM, Sha Z, Williams DA, Pletnev AA, de Bruin G, et al. Inhibition of the Proteasome beta2 Site Sensitizes Triple-Negative Breast Cancer Cells to beta5 Inhibitors and Suppresses Nrf1 Activation. Cell Chem Biol. 2017;24(2):218–30.

46. Radhakrishnan SK, Lee CS, Young P, Beskow A, Chan JY, Deshaies RJ. Transcription factor Nrf1 mediates the proteasome recovery pathway after proteasome inhibition in mammalian cells. Mol Cell. 2010;38(1):17–28.

47. Lochmann TL, Powell KM, Ham J, Floros KV, Heisey DAR, Kurupi RIJ, et al. Targeted inhibition of histone H3K27 demethylation is effective in high-risk neuroblastoma. Sci Transl Med. 2018;10(441).

48. Floros KV, Lochmann TL, Hu B, Monterrubio C, Hughes MT, Wells JD, et al. Coamplification of miR-4728 protects HER2-amplified breast cancers from targeted therapy. Proceedings of the National Academy of Sciences of the United States of America. 2018;115(11):E2594–E603.

49. Turner TH, Alzubi MA, Sohal SS, Olex AL, Dozmorov MG, Harrell JC. Characterizing the efficacy of cancer therapeutics in patient-derived xenograft models of metastatic breast cancer. Breast Cancer Res Treat. 2018;170(2):221–34.

50. Hampton JD, Peterson EJ, Katner SJ, Turner TH, Alzubi MA, Harrell JC, et al. Exploitation of Sulfated Glycosaminoglycan Status for Precision Medicine of Triplatin in Triple-Negative Breast Cancer. Mol Cancer Ther. 2022;21(2):271–81.

