## Supplementary figures and images for "Genomic screening reveals UBA1 as a potent and druggable target in c-MYC-high TNBC models"

### Supplemental Figure S1

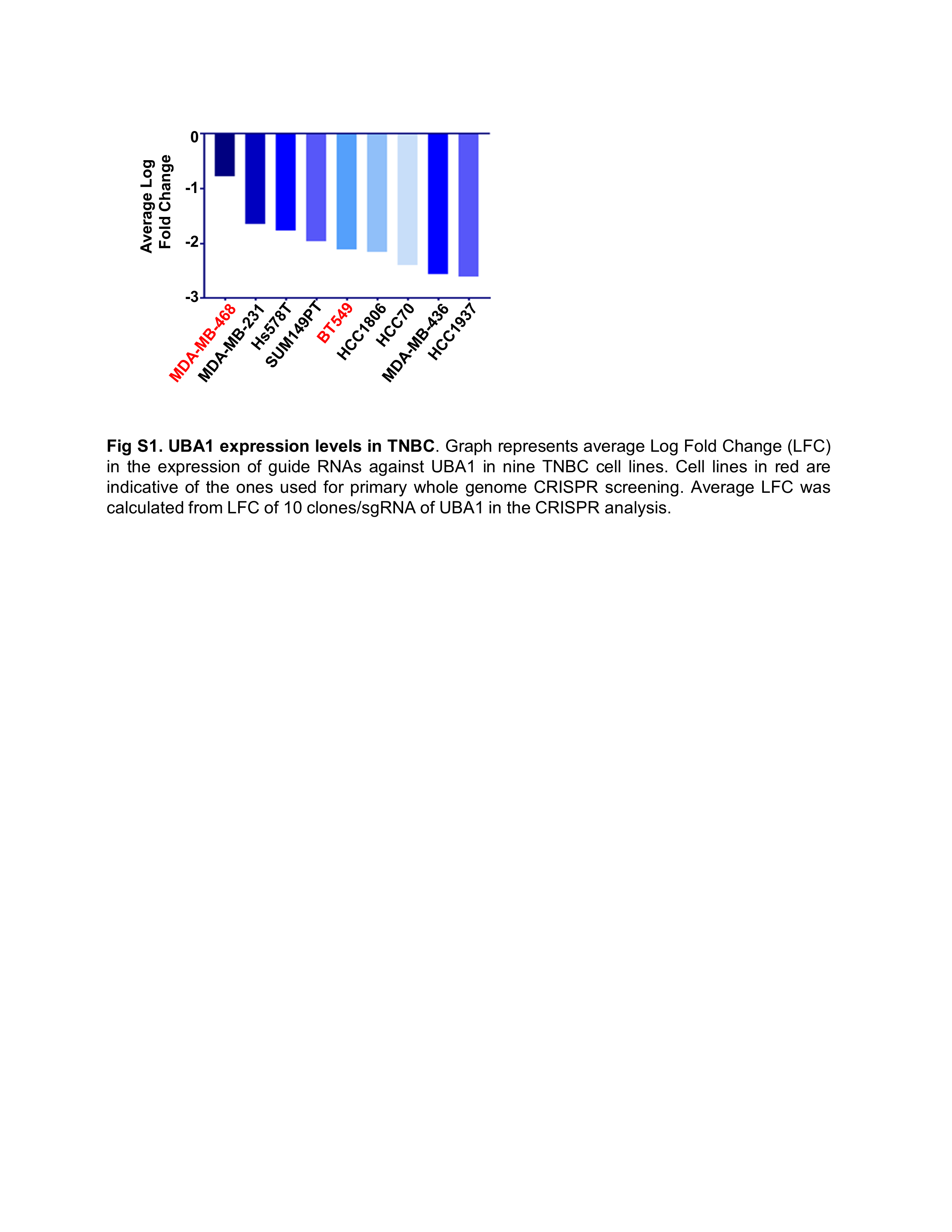

### Supplemental Figure S2

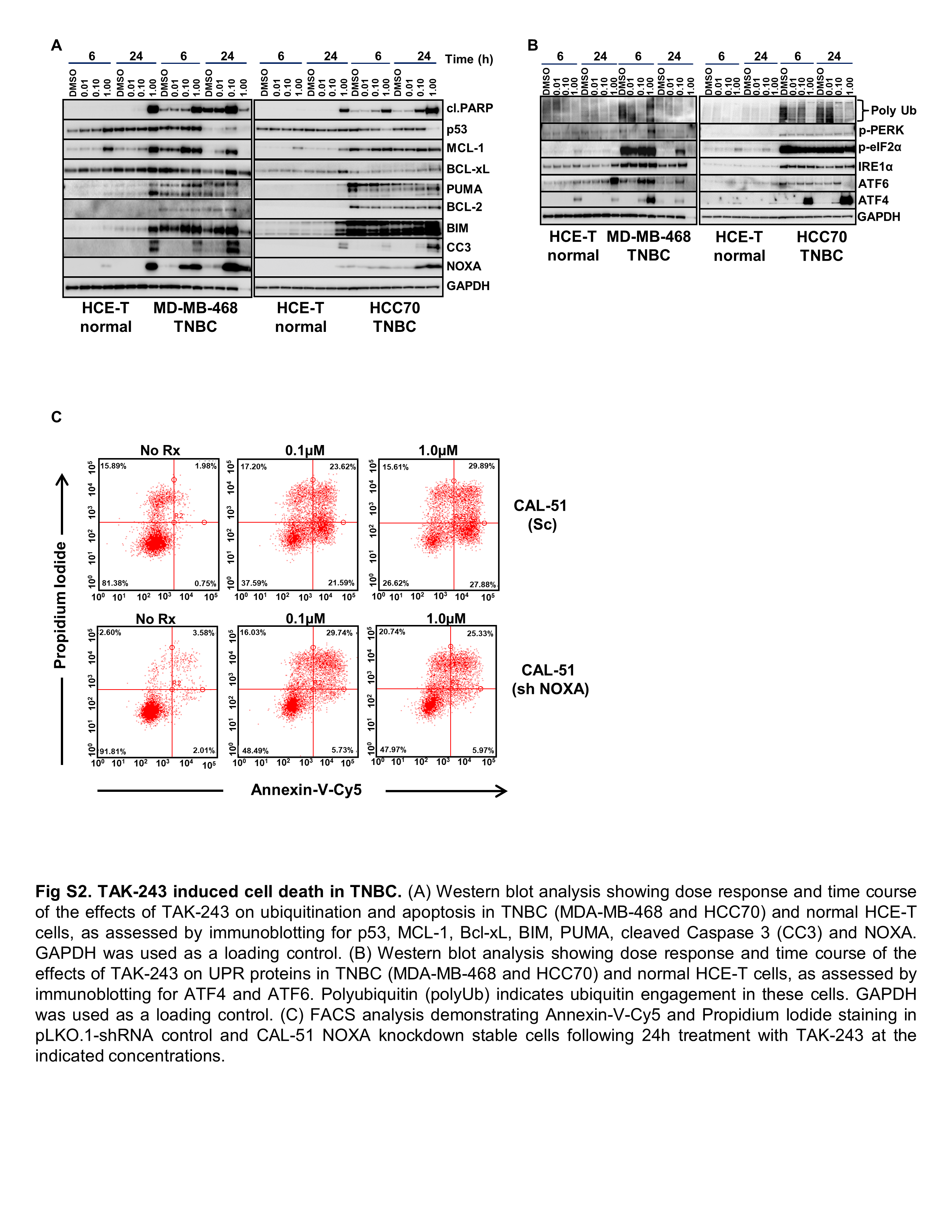

### Supplemental Figure S3

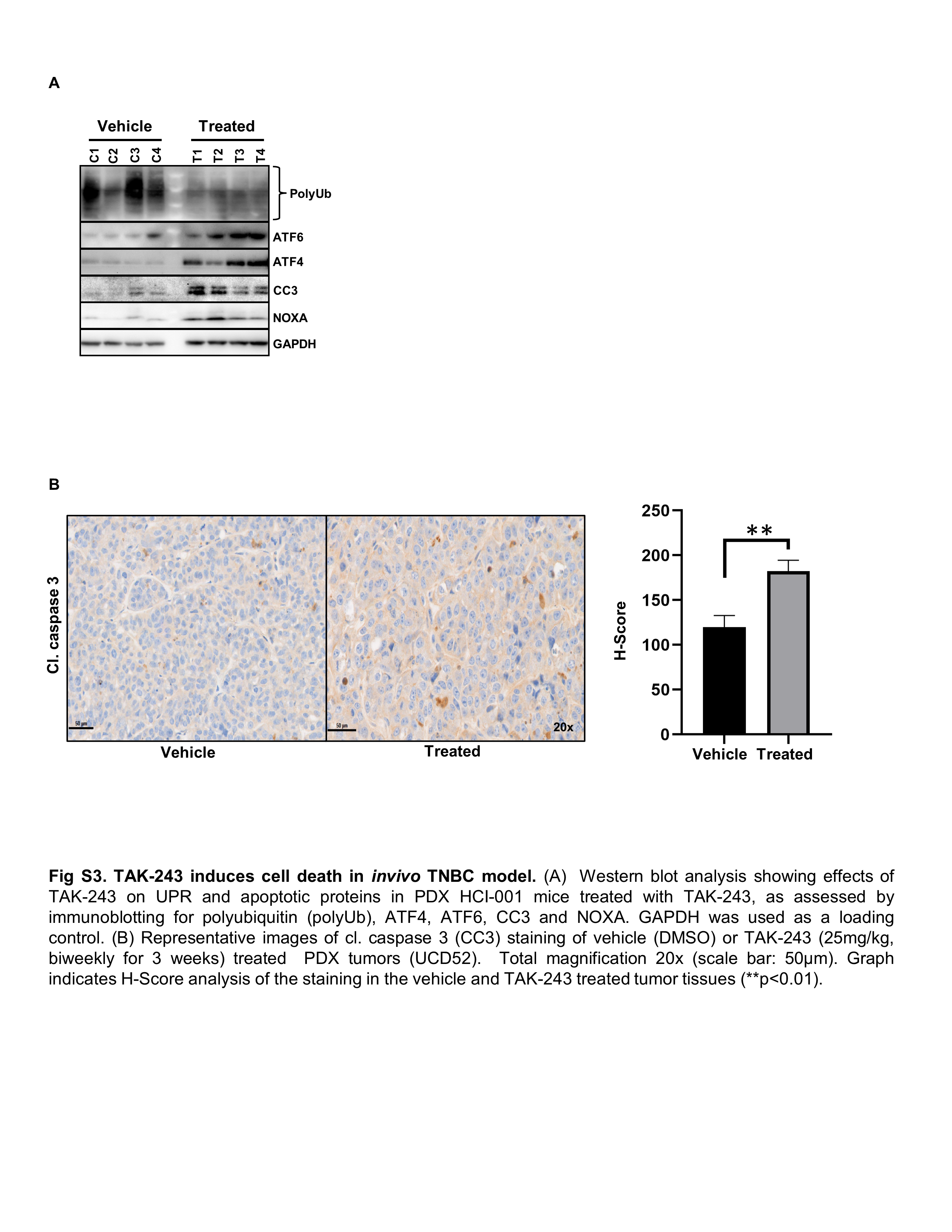

### Supplemental Figure S4

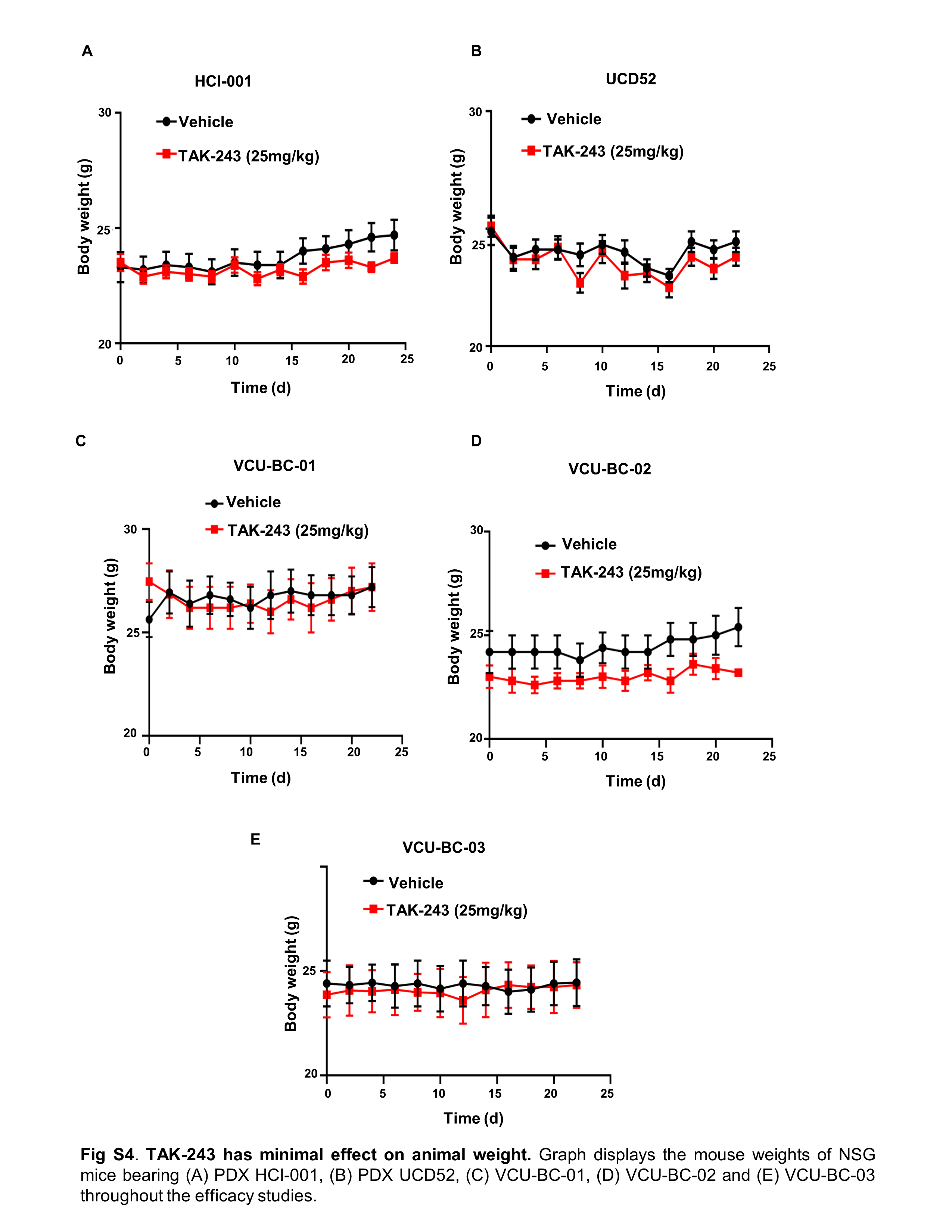

### Supplemental Figure S5

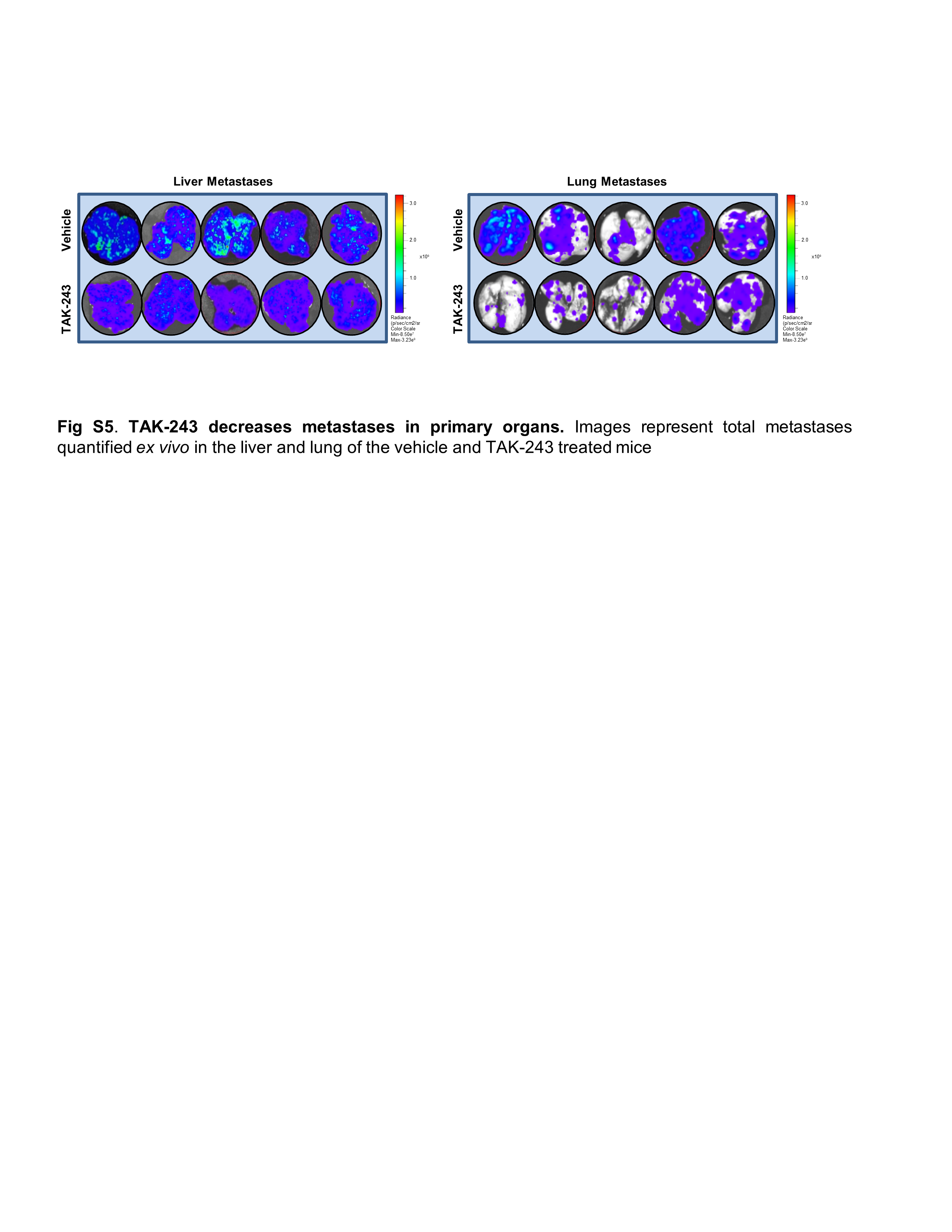

### Supplemental table T2

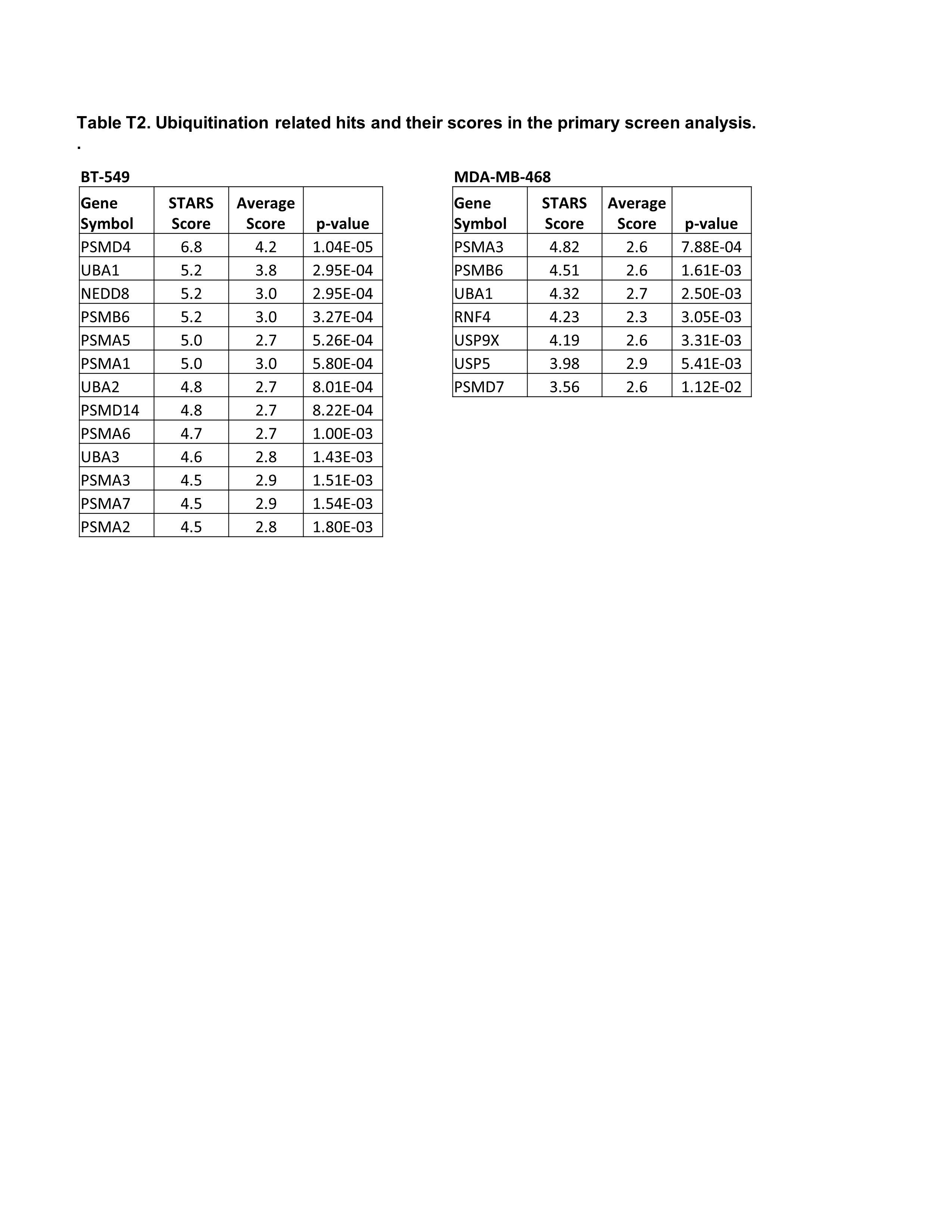

### Supplemental table T3

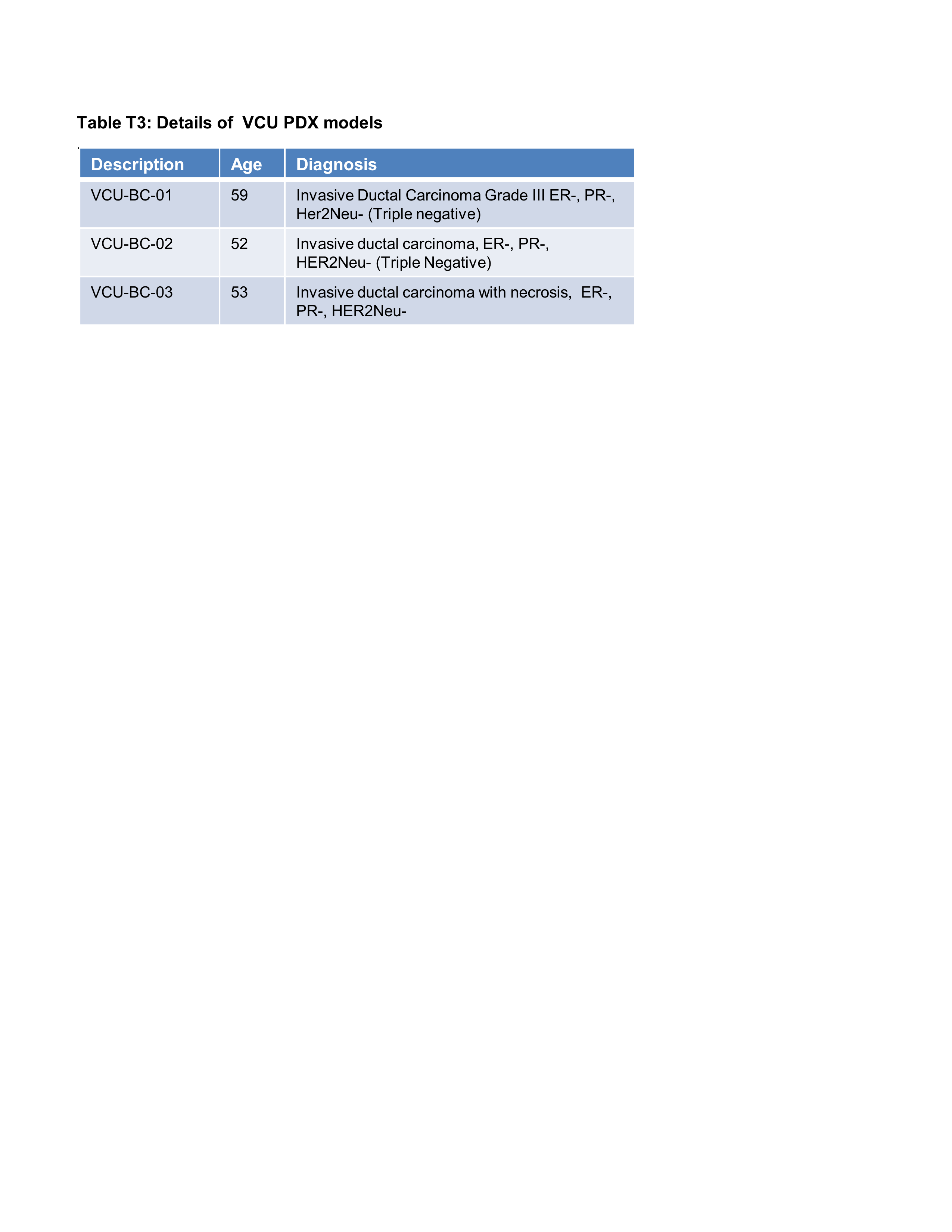
