## Supplemental table T1 for "Genomic screening reveals UBA1 as a potent and druggable target in c-MYC-high TNBC models"

**Table 1.** Top 400 genes (approximately 2% of total genes targeted) by STARS scores.

| **BT549** | | | |
|---|---|---|---|
| Gene Symbol | STARS Score | Average Score | p-value |
| RAD9A | 8.25 | 5.3 | 1.04E-06 |
| RNGTT | 8.1 | 5.3 | 1.04E-06 |
| SBDS | 7.62 | 5.41 | 2.08E-06 |
| PRC1 | 7.5 | 4.81 | 2.08E-06 |
| BUB3 | 7.4 | 4.75 | 3.12E-06 |
| C1orf109 | 7.29 | 4.73 | 4.16E-06 |
| PDCD2 | 7.27 | 4.57 | 4.16E-06 |
| LSM2 | 7.18 | 4.25 | 4.16E-06 |
| ATP6V0C | 7.1 | 4.71 | 4.16E-06 |
| GINS2 | 7.08 | 4.67 | 4.16E-06 |
| ATP6V1F | 7.05 | 4.42 | 4.16E-06 |
| RFC3 | 7.02 | 4.12 | 4.16E-06 |
| TXNL4A | 6.99 | 4.59 | 5.20E-06 |
| WDR82 | 6.95 | 4.21 | 6.24E-06 |
| HSD17B12 | 6.92 | 4.58 | 7.28E-06 |
| CHAF1B | 6.91 | 4.67 | 7.28E-06 |
| MCM5 | 6.9 | 4 | 7.28E-06 |
| ACTR1A | 6.87 | 4.57 | 8.32E-06 |
| TINF2 | 6.79 | 3.97 | 1.04E-05 |
| ELP5 | 6.77 | 4.02 | 1.04E-05 |
| PSMD4 | 6.76 | 4.17 | 1.04E-05 |
| CIRH1A | 6.71 | 4.13 | 1.14E-05 |
| RCC1 | 6.7 | 4.38 | 1.14E-05 |
| MRPL33 | 6.64 | 4.13 | 1.35E-05 |
| MAD2L2 | 6.63 | 4.01 | 1.35E-05 |
| GAPDH | 6.62 | 4.07 | 1.35E-05 |
| YARS | 6.6 | 3.93 | 1.35E-05 |
| GTF2H4 | 6.58 | 3.94 | 1.35E-05 |
| EXOSC6 | 6.58 | 4.71 | 1.46E-05 |
| VPS25 | 6.5 | 4.79 | 1.56E-05 |
| NAPG | 6.41 | 3.65 | 2.08E-05 |
| SS18L2 | 6.4 | 4.45 | 2.08E-05 |
| MARS2 | 6.38 | 4.43 | 2.18E-05 |
| RPL12 | 6.35 | 3.82 | 2.29E-05 |
| LONP1 | 6.35 | 4.04 | 2.29E-05 |
| FDXR | 6.33 | 4.09 | 2.49E-05 |
| DHX37 | 6.32 | 3.76 | 2.60E-05 |
| NCAPG | 6.3 | 4.22 | 2.70E-05 |
| POLR2L | 6.27 | 3.98 | 3.01E-05 |
| SOD1 | 6.27 | 3.98 | 3.01E-05 |
| ATP1A1 | 6.25 | 3.64 | 3.12E-05 |
| YARS2 | 6.24 | 5.18 | 3.12E-05 |

| **MDA-MB-468** | | | |
|---|---|---|---|
| Gene Symbol | STARS Score | Average Score | p-value |
| NIP7 | 8.31 | 5.14 | 1.04E-06 |
| C1orf109 | 8.24 | 4.98 | 1.04E-06 |
| WDR82 | 8.23 | 4.89 | 1.04E-06 |
| MRPS11 | 8.01 | 5.28 | 1.04E-06 |
| WDR18 | 7.75 | 4.84 | 2.08E-06 |
| MARS2 | 7.75 | 4.89 | 2.08E-06 |
| TAMM41 | 7.54 | 5.13 | 2.08E-06 |
| CRCP | 7.51 | 5.09 | 2.08E-06 |
| NARFL | 7.5 | 5.03 | 2.08E-06 |
| DYNLRB1 | 7.4 | 5 | 3.12E-06 |
| FARSB | 7.09 | 4.15 | 4.16E-06 |
| DBR1 | 6.99 | 4.99 | 5.20E-06 |
| LONP1 | 6.82 | 4.47 | 1.04E-05 |
| ZBTB11 | 6.81 | 4.05 | 1.04E-05 |
| SPATA5L1 | 6.71 | 4.79 | 1.14E-05 |
| ATP6V1B2 | 6.68 | 3.82 | 1.25E-05 |
| ANKLE2 | 6.49 | 4.03 | 1.56E-05 |
| SS18L2 | 6.43 | 4.64 | 1.77E-05 |
| PMPCB | 6.42 | 3.98 | 1.98E-05 |
| SOD2 | 6.41 | 3.84 | 1.98E-05 |
| ZMAT2 | 6.3 | 3.6 | 2.60E-05 |
| SCAP | 6.29 | 4.39 | 2.81E-05 |
| YARS2 | 6.28 | 4.04 | 2.91E-05 |
| MMS22L | 6.26 | 3.55 | 3.12E-05 |
| SARS | 6.25 | 3.7 | 3.12E-05 |
| TAF6 | 6.14 | 4.46 | 3.64E-05 |
| EXOSC8 | 6.07 | 3.59 | 4.26E-05 |
| PPP1R8 | 6.03 | 3.88 | 4.78E-05 |
| POLR2L | 6.02 | 3.93 | 4.89E-05 |
| LETM1 | 6.01 | 4.41 | 4.89E-05 |
| MAD2L2 | 5.99 | 3.62 | 5.20E-05 |
| CDAN1 | 5.96 | 3.71 | 5.72E-05 |
| CYC1 | 5.93 | 3.68 | 5.82E-05 |
| BUB3 | 5.92 | 4.34 | 5.82E-05 |
| TAF1C | 5.91 | 3.74 | 5.93E-05 |
| HSPE1 | 5.9 | 3.36 | 6.13E-05 |
| SNAPC5 | 5.89 | 3.71 | 6.44E-05 |
| RPL23 | 5.88 | 3.64 | 6.44E-05 |
| NOA1 | 5.87 | 3.53 | 6.76E-05 |
| CIRH1A | 5.85 | 3.88 | 6.76E-05 |
| DPAGT1 | 5.85 | 4.37 | 6.96E-05 |
| PELP1 | 5.77 | 3.47 | 7.80E-05 |

| | | | |
|---|---|---|---|
| TOP2A | 6.21 | 3.68 | 3.43E-05 |
| C9orf114 | 6.18 | 3.69 | 3.64E-05 |
| RFC2 | 6.17 | 5.1 | 3.64E-05 |
| TPI1 | 6.16 | 3.53 | 3.64E-05 |
| POLR3A | 6.16 | 3.97 | 3.64E-05 |
| DYNLRB1 | 6.16 | 4.02 | 3.64E-05 |
| NOP58 | 6.15 | 3.96 | 3.64E-05 |
| WDR55 | 6.06 | 4.01 | 4.26E-05 |
| TAF1C | 6 | 4.13 | 5.20E-05 |
| DYNC1I2 | 6 | 3.8 | 5.20E-05 |
| MNAT1 | 5.98 | 3.44 | 5.30E-05 |
| ZMAT2 | 5.98 | 3.91 | 5.41E-05 |
| GPN1 | 5.97 | 3.85 | 5.61E-05 |
| RPL4 | 5.94 | 3.94 | 5.82E-05 |
| DPAGT1 | 5.93 | 4.07 | 5.82E-05 |
| MARS | 5.89 | 3.87 | 6.34E-05 |
| ESPL1 | 5.87 | 4.24 | 6.44E-05 |
| RPL37 | 5.87 | 3.64 | 6.65E-05 |
| RNMT | 5.86 | 3.7 | 6.76E-05 |
| NELFB | 5.86 | 4.22 | 6.76E-05 |
| NHLRC2 | 5.86 | 3.4 | 6.76E-05 |
| MRPS11 | 5.85 | 3.71 | 6.76E-05 |
| GUK1 | 5.85 | 4.35 | 6.76E-05 |
| CPSF4 | 5.85 | 3.32 | 6.96E-05 |
| GGPS1 | 5.84 | 3.68 | 6.96E-05 |
| PRELID1 | 5.83 | 4.03 | 6.96E-05 |
| TARDBP | 5.81 | 3.38 | 7.07E-05 |
| PWP2 | 5.81 | 3.55 | 7.28E-05 |
| OR4F21 | 5.8 | 3.41 | 7.48E-05 |
| WDR18 | 5.79 | 3.83 | 7.59E-05 |
| LUC7L3 | 5.77 | 3.66 | 7.80E-05 |
| GINS1 | 5.76 | 4.26 | 8.11E-05 |
| GTPBP4 | 5.75 | 4.34 | 8.21E-05 |
| CDC123 | 5.75 | 4.3 | 8.52E-05 |
| RPS3A | 5.74 | 3.62 | 8.52E-05 |
| TRAPPC3 | 5.74 | 3.27 | 8.52E-05 |
| RPL7 | 5.73 | 3.18 | 8.94E-05 |
| SARS | 5.72 | 3.84 | 8.94E-05 |
| NCBP2 | 5.72 | 4.09 | 9.15E-05 |
| RGPD8 | 5.71 | 3.11 | 9.25E-05 |
| BIRC5 | 5.7 | 3.7 | 9.67E-05 |
| ARL2 | 5.69 | 3.37 | 9.88E-05 |
| NOL10 | 5.69 | 3.76 | 9.88E-05 |
| HSPE1-MOB4 | 5.68 | 3.42 | 9.88E-05 |
| LRPPRC | 5.68 | 4.06 | 1.03E-04 |
| EEF2 | 5.66 | 3.78 | 1.06E-04 |
| EIF2B1 | 5.65 | 3.48 | 1.09E-04 |

| | | | |
|---|---|---|---|
| GEMIN5 | 5.76 | 3.68 | 7.80E-05 |
| ALG2 | 5.74 | 3.64 | 8.52E-05 |
| PPP1R2 | 5.71 | 4.1 | 9.15E-05 |
| RFC3 | 5.71 | 3.82 | 9.36E-05 |
| GTF3A | 5.71 | 3.77 | 9.36E-05 |
| COPS6 | 5.69 | 3.96 | 9.88E-05 |
| RPP40 | 5.67 | 3.44 | 1.03E-04 |
| THOC3 | 5.64 | 3.39 | 1.11E-04 |
| COQ4 | 5.63 | 3.25 | 1.11E-04 |
| PMPCA | 5.63 | 3.61 | 1.12E-04 |
| NSF | 5.62 | 3.64 | 1.19E-04 |
| DNAJC9 | 5.6 | 3.51 | 1.26E-04 |
| SOD1 | 5.6 | 3.81 | 1.26E-04 |
| FXN | 5.59 | 3.97 | 1.27E-04 |
| EARS2 | 5.55 | 3.5 | 1.36E-04 |
| VPS25 | 5.55 | 3.97 | 1.37E-04 |
| SMG6 | 5.53 | 3.11 | 1.46E-04 |
| FBL | 5.51 | 3.71 | 1.53E-04 |
| TXNL4A | 5.5 | 3.58 | 1.56E-04 |
| ACTR1A | 5.47 | 3.54 | 1.65E-04 |
| MRPL18 | 5.47 | 3.14 | 1.66E-04 |
| MVK | 5.47 | 3.46 | 1.66E-04 |
| RPS4X | 5.45 | 3.48 | 1.70E-04 |
| TBC1D3C | 5.44 | 3.25 | 1.77E-04 |
| ATP6V1F | 5.42 | 3.23 | 1.83E-04 |
| SNAPC1 | 5.39 | 3.22 | 1.91E-04 |
| CPSF4 | 5.38 | 3.42 | 1.98E-04 |
| NDUFAF3 | 5.37 | 3.62 | 2.00E-04 |
| TBC1D3H | 5.37 | 3.23 | 2.00E-04 |
| SRSF2 | 5.37 | 3.27 | 2.05E-04 |
| NDUFA8 | 5.34 | 3.56 | 2.23E-04 |
| NELFB | 5.33 | 3.91 | 2.30E-04 |
| COX15 | 5.33 | 3.27 | 2.30E-04 |
| CDC123 | 5.33 | 3.32 | 2.32E-04 |
| RAE1 | 5.33 | 2.93 | 2.34E-04 |
| C1QBP | 5.31 | 3.15 | 2.48E-04 |
| SF3B5 | 5.29 | 3.23 | 2.59E-04 |
| FNTB | 5.29 | 3.56 | 2.61E-04 |
| TBC1D3G | 5.28 | 3.04 | 2.65E-04 |
| TBC1D3 | 5.28 | 3.04 | 2.65E-04 |
| RNPC3 | 5.27 | 3.63 | 2.74E-04 |
| FARS2 | 5.26 | 3.51 | 2.78E-04 |
| INTS3 | 5.23 | 3.6 | 2.95E-04 |
| ROMO1 | 5.22 | 3.57 | 2.98E-04 |
| NDUFB4 | 5.22 | 3.1 | 3.04E-04 |
| CHMP6 | 5.21 | 3.76 | 3.09E-04 |
| TUBGCP2 | 5.18 | 3.47 | 3.28E-04 |

| | | | |
|---|---|---|---|
| GEMIN5 | 5.64 | 3.35 | 1.10E-04 |
| DDX54 | 5.63 | 3.32 | 1.12E-04 |
| TARS | 5.6 | 3.61 | 1.23E-04 |
| CCT5 | 5.58 | 3.53 | 1.29E-04 |
| SRP14 | 5.58 | 3.99 | 1.29E-04 |
| CRCP | 5.58 | 3.45 | 1.30E-04 |
| PMPCA | 5.56 | 3.25 | 1.34E-04 |
| POLR1C | 5.55 | 4.63 | 1.35E-04 |
| POLD3 | 5.55 | 3.13 | 1.35E-04 |
| MRPL10 | 5.52 | 3.36 | 1.48E-04 |
| CDC23 | 5.52 | 3.51 | 1.50E-04 |
| DNAJC17 | 5.5 | 3.96 | 1.56E-04 |
| PHB2 | 5.49 | 3.58 | 1.57E-04 |
| MMS22L | 5.48 | 3.5 | 1.59E-04 |
| NPIPA5 | 5.48 | 3.42 | 1.59E-04 |
| WDHD1 | 5.47 | 3.19 | 1.64E-04 |
| RPL17-C18orf3 | 5.46 | 3.24 | 1.66E-04 |
| CHMP6 | 5.44 | 3.77 | 1.77E-04 |
| MCM6 | 5.43 | 3.36 | 1.80E-04 |
| FBL | 5.43 | 3.61 | 1.80E-04 |
| SARS2 | 5.43 | 3.02 | 1.82E-04 |
| AFG3L2 | 5.42 | 3.41 | 1.83E-04 |
| MRPL16 | 5.42 | 3.26 | 1.83E-04 |
| DCTN5 | 5.42 | 3.25 | 1.83E-04 |
| RSL1D1 | 5.38 | 3.21 | 1.98E-04 |
| AARS | 5.38 | 3.41 | 1.99E-04 |
| POLR1A | 5.37 | 3.2 | 2.00E-04 |
| ALG13 | 5.37 | 3.86 | 2.00E-04 |
| METTL16 | 5.36 | 3.16 | 2.06E-04 |
| SRSF2 | 5.36 | 3.36 | 2.11E-04 |
| CWC22 | 5.36 | 3 | 2.11E-04 |
| DBR1 | 5.36 | 4.21 | 2.11E-04 |
| EARS2 | 5.35 | 3.59 | 2.14E-04 |
| POLD1 | 5.35 | 3.04 | 2.20E-04 |
| HCFC1 | 5.33 | 3.77 | 2.31E-04 |
| SOD2 | 5.31 | 3.09 | 2.48E-04 |
| TRNT1 | 5.31 | 4.01 | 2.48E-04 |
| SAE1 | 5.3 | 3.35 | 2.51E-04 |
| GINS3 | 5.29 | 3.83 | 2.59E-04 |
| GFM1 | 5.29 | 3.31 | 2.61E-04 |
| ALDOA | 5.29 | 3.11 | 2.62E-04 |
| NSF | 5.27 | 3.21 | 2.68E-04 |
| COASY | 5.26 | 3.28 | 2.79E-04 |
| PHF5A | 5.26 | 3.04 | 2.84E-04 |
| XRCC6 | 5.25 | 3.31 | 2.87E-04 |
| POLR3H | 5.25 | 4.33 | 2.91E-04 |
| UBA1 | 5.24 | 3.78 | 2.94E-04 |

| | | | |
|---|---|---|---|
| PAXBP1 | 5.17 | 2.94 | 3.36E-04 |
| LARS2 | 5.16 | 3.14 | 3.43E-04 |
| POLR3A | 5.15 | 3.12 | 3.58E-04 |
| CDK7 | 5.14 | 3.44 | 3.63E-04 |
| COG3 | 5.14 | 3.17 | 3.69E-04 |
| GINS2 | 5.14 | 3.3 | 3.72E-04 |
| PRKRA | 5.13 | 3.6 | 3.77E-04 |
| DDX49 | 5.09 | 3.74 | 4.13E-04 |
| TIMM10 | 5.08 | 3.44 | 4.20E-04 |
| MRPL57 | 5.07 | 3.34 | 4.25E-04 |
| NDNL2 | 5.06 | 3.37 | 4.41E-04 |
| RBM4 | 5.05 | 3.11 | 4.58E-04 |
| HARS2 | 5.03 | 2.83 | 4.89E-04 |
| CCDC115 | 5.02 | 3.36 | 4.95E-04 |
| PET117 | 5.01 | 3.81 | 5.05E-04 |
| MRPL33 | 4.98 | 3.51 | 5.42E-04 |
| RPL9 | 4.94 | 3.5 | 5.95E-04 |
| PDCD2 | 4.94 | 3.58 | 6.01E-04 |
| RPAIN | 4.94 | 3.06 | 6.01E-04 |
| RBM25 | 4.92 | 3.01 | 6.38E-04 |
| LUC7L3 | 4.89 | 3.31 | 6.83E-04 |
| EEF2 | 4.89 | 4.04 | 6.86E-04 |
| RPS15A | 4.89 | 3.38 | 6.86E-04 |
| COPG1 | 4.89 | 3.37 | 6.87E-04 |
| ATP5E | 4.89 | 2.98 | 6.93E-04 |
| SNAPC2 | 4.89 | 3.43 | 6.93E-04 |
| CCT3 | 4.88 | 2.77 | 6.95E-04 |
| PGK1 | 4.88 | 3.44 | 6.95E-04 |
| MYC | 4.88 | 2.95 | 7.02E-04 |
| MRPL38 | 4.88 | 3.7 | 7.02E-04 |
| UTP15 | 4.87 | 2.9 | 7.05E-04 |
| EEF1A1 | 4.85 | 3.15 | 7.35E-04 |
| PSMG4 | 4.85 | 3.35 | 7.43E-04 |
| HYOU1 | 4.84 | 3.38 | 7.59E-04 |
| VBP1 | 4.84 | 3.48 | 7.61E-04 |
| RIC1 | 4.84 | 2.73 | 7.63E-04 |
| CCDC86 | 4.83 | 2.71 | 7.73E-04 |
| CDC7 | 4.83 | 2.79 | 7.73E-04 |
| TRAPPC8 | 4.83 | 3.65 | 7.77E-04 |
| PSMA3 | 4.82 | 2.62 | 7.88E-04 |
| RPS28 | 4.82 | 3.76 | 7.89E-04 |
| PFDN2 | 4.82 | 3.03 | 8.01E-04 |
| RAD9A | 4.77 | 3.07 | 9.00E-04 |
| POLR1C | 4.76 | 3.52 | 9.15E-04 |
| PDCD11 | 4.76 | 3.31 | 9.18E-04 |
| DNAJC17 | 4.74 | 3.35 | 9.55E-04 |
| RPL19 | 4.73 | 2.92 | 9.78E-04 |

| | | | |
|---|---|---|---|
| LETM1 | 5.24 | 3.82 | 2.94E-04 |
| SF3A1 | 5.23 | 3.52 | 2.94E-04 |
| RPTOR | 5.23 | 3.27 | 2.95E-04 |
| NEDD8 | 5.23 | 3.03 | 2.95E-04 |
| EXOSC8 | 5.22 | 3.19 | 3.04E-04 |
| SDHB | 5.22 | 3.38 | 3.04E-04 |
| UBE2N | 5.22 | 3.65 | 3.04E-04 |
| KIF18A | 5.21 | 3.51 | 3.04E-04 |
| RABGGTB | 5.2 | 3.4 | 3.11E-04 |
| MRPL28 | 5.2 | 3.1 | 3.11E-04 |
| CIAO1 | 5.2 | 3.64 | 3.20E-04 |
| TONSL | 5.19 | 3.71 | 3.21E-04 |
| MLST8 | 5.19 | 3.05 | 3.27E-04 |
| PSMB6 | 5.19 | 2.96 | 3.27E-04 |
| MVK | 5.18 | 3.27 | 3.28E-04 |
| IPO9 | 5.18 | 3.19 | 3.30E-04 |
| CDC45 | 5.18 | 2.93 | 3.33E-04 |
| DDX55 | 5.17 | 3.7 | 3.35E-04 |
| SNRNP25 | 5.17 | 3.76 | 3.35E-04 |
| PSMG4 | 5.17 | 3.52 | 3.35E-04 |
| FCF1 | 5.17 | 3.36 | 3.41E-04 |
| TTI1 | 5.16 | 2.89 | 3.42E-04 |
| RUVBL1 | 5.16 | 3.76 | 3.43E-04 |
| EIF1AX | 5.15 | 3.7 | 3.62E-04 |
| ORAOV1 | 5.14 | 3.72 | 3.67E-04 |
| UBE2L3 | 5.14 | 3.19 | 3.69E-04 |
| CCT3 | 5.13 | 3.12 | 3.77E-04 |
| URB1 | 5.13 | 3.17 | 3.77E-04 |
| RBM22 | 5.13 | 2.78 | 3.84E-04 |
| MAK16 | 5.12 | 3.4 | 3.89E-04 |
| POLRMT | 5.12 | 3.07 | 3.89E-04 |
| NOP2 | 5.12 | 3.07 | 3.94E-04 |
| HDAC3 | 5.11 | 3.2 | 3.96E-04 |
| RPAP1 | 5.11 | 3.73 | 3.96E-04 |
| OSGEP | 5.11 | 3.45 | 3.97E-04 |
| TRMT112 | 5.1 | 3.7 | 4.02E-04 |
| ANKLE2 | 5.1 | 3.48 | 4.02E-04 |
| CCDC84 | 5.09 | 3.2 | 4.15E-04 |
| EIF3I | 5.08 | 2.85 | 4.20E-04 |
| TEN1 | 5.06 | 3.19 | 4.41E-04 |
| PGK1 | 5.06 | 3.11 | 4.51E-04 |
| RPL31 | 5.04 | 3.12 | 4.66E-04 |
| LYRM4 | 5.04 | 3.37 | 4.78E-04 |
| LARS2 | 5.03 | 3.16 | 4.88E-04 |
| SNAPC5 | 5.03 | 3.2 | 4.92E-04 |
| RPL34 | 5.02 | 2.99 | 5.02E-04 |
| PKM | 5.02 | 3.23 | 5.05E-04 |

| | | | |
|---|---|---|---|
| RPL35 | 4.72 | 2.92 | 9.86E-04 |
| CENPW | 4.71 | 2.94 | 1.01E-03 |
| FAM96B | 4.71 | 3.25 | 1.02E-03 |
| DHX33 | 4.71 | 3.39 | 1.02E-03 |
| HEATR1 | 4.7 | 2.99 | 1.03E-03 |
| MTPAP | 4.7 | 3.08 | 1.03E-03 |
| HNRNPM | 4.69 | 3.66 | 1.06E-03 |
| XRCC3 | 4.69 | 3.3 | 1.07E-03 |
| YARS | 4.68 | 2.8 | 1.07E-03 |
| RNGTT | 4.68 | 3.26 | 1.07E-03 |
| POP5 | 4.68 | 3.53 | 1.08E-03 |
| DHX37 | 4.67 | 3.13 | 1.11E-03 |
| SAE1 | 4.67 | 3.4 | 1.11E-03 |
| NDUFB11 | 4.66 | 2.94 | 1.14E-03 |
| CAP1 | 4.65 | 3.3 | 1.16E-03 |
| MIPEP | 4.65 | 3.01 | 1.16E-03 |
| NDUFB10 | 4.65 | 3.23 | 1.16E-03 |
| NDUFAF1 | 4.64 | 3.04 | 1.19E-03 |
| CHAF1B | 4.64 | 3.2 | 1.20E-03 |
| SBDS | 4.63 | 3.56 | 1.21E-03 |
| NOP9 | 4.63 | 3.6 | 1.22E-03 |
| SMU1 | 4.62 | 2.94 | 1.24E-03 |
| CYCS | 4.61 | 3.11 | 1.29E-03 |
| UQCRC2 | 4.61 | 3.05 | 1.29E-03 |
| MCM4 | 4.61 | 2.96 | 1.29E-03 |
| SRBD1 | 4.6 | 2.74 | 1.30E-03 |
| RUVBL2 | 4.6 | 2.71 | 1.31E-03 |
| HSPE1-MOB4 | 4.6 | 2.98 | 1.31E-03 |
| CTCF | 4.6 | 3.66 | 1.32E-03 |
| EMC3 | 4.56 | 2.75 | 1.43E-03 |
| POLR3H | 4.56 | 3.71 | 1.44E-03 |
| MRPS23 | 4.55 | 3.03 | 1.46E-03 |
| TKT | 4.55 | 2.77 | 1.47E-03 |
| DCTN5 | 4.54 | 2.8 | 1.48E-03 |
| TUBB | 4.54 | 2.91 | 1.48E-03 |
| C21orf59 | 4.52 | 3.2 | 1.57E-03 |
| BOP1 | 4.52 | 3.12 | 1.59E-03 |
| POLD3 | 4.51 | 2.65 | 1.60E-03 |
| C3orf17 | 4.51 | 3.16 | 1.60E-03 |
| H2AFX | 4.51 | 3.16 | 1.61E-03 |
| PSMB6 | 4.51 | 2.57 | 1.62E-03 |
| NDUFB6 | 4.5 | 3.8 | 1.66E-03 |
| LRR1 | 4.49 | 3.08 | 1.67E-03 |
| GNL3 | 4.49 | 3.11 | 1.69E-03 |
| GINS1 | 4.49 | 3.33 | 1.69E-03 |
| BRIX1 | 4.48 | 2.8 | 1.72E-03 |
| DTYMK | 4.48 | 3.37 | 1.72E-03 |

| | | | |
|---|---|---|---|
| MCM3 | 5.01 | 3.78 | 5.05E-04 |
| CYCS | 5.01 | 2.9 | 5.14E-04 |
| DDX10 | 5.01 | 3.39 | 5.15E-04 |
| ATP6V1B2 | 5.01 | 3.8 | 5.18E-04 |
| SFSWAP | 5 | 3.33 | 5.24E-04 |
| PSMA5 | 4.99 | 2.73 | 5.26E-04 |
| PRMT1 | 4.99 | 3.68 | 5.28E-04 |
| SF3A2 | 4.97 | 2.84 | 5.47E-04 |
| RABGGTA | 4.97 | 3.54 | 5.48E-04 |
| BUB1B | 4.97 | 2.93 | 5.49E-04 |
| PPA1 | 4.96 | 3.29 | 5.76E-04 |
| PSMA1 | 4.95 | 3.03 | 5.80E-04 |
| MYC | 4.95 | 2.9 | 5.83E-04 |
| MRPL21 | 4.95 | 2.96 | 5.93E-04 |
| RPL23 | 4.95 | 3.15 | 5.94E-04 |
| C3orf17 | 4.94 | 2.85 | 5.99E-04 |
| NPLOC4 | 4.94 | 3.27 | 6.07E-04 |
| GTF2A2 | 4.92 | 3.36 | 6.26E-04 |
| INTS3 | 4.92 | 3.72 | 6.30E-04 |
| SNRPG | 4.91 | 3.38 | 6.39E-04 |
| CDCA8 | 4.91 | 3.16 | 6.39E-04 |
| FARSB | 4.9 | 4.43 | 6.66E-04 |
| MRPS23 | 4.9 | 3.52 | 6.66E-04 |
| SPATA5L1 | 4.9 | 3.19 | 6.74E-04 |
| SF1 | 4.9 | 2.9 | 6.76E-04 |
| MRPL39 | 4.89 | 3.53 | 6.87E-04 |
| THOC3 | 4.89 | 3.31 | 6.93E-04 |
| RBBP5 | 4.88 | 3.27 | 6.95E-04 |
| DDB1 | 4.88 | 3.05 | 6.95E-04 |
| RPL14 | 4.88 | 3.17 | 7.03E-04 |
| RPL24 | 4.88 | 2.89 | 7.04E-04 |
| E2F3 | 4.88 | 2.91 | 7.04E-04 |
| PIK3C3 | 4.87 | 2.85 | 7.06E-04 |
| DHPS | 4.87 | 3.06 | 7.16E-04 |
| CCT2 | 4.86 | 3.24 | 7.22E-04 |
| METTL14 | 4.85 | 2.91 | 7.34E-04 |
| NVL | 4.85 | 3.28 | 7.43E-04 |
| CCT4 | 4.85 | 2.95 | 7.44E-04 |
| TOP3A | 4.84 | 2.87 | 7.45E-04 |
| RPL27A | 4.84 | 3.27 | 7.60E-04 |
| NMD3 | 4.84 | 3.05 | 7.61E-04 |
| VCP | 4.83 | 2.85 | 7.71E-04 |
| IARS2 | 4.83 | 4.04 | 7.73E-04 |
| CENPN | 4.83 | 2.75 | 7.73E-04 |
| OBFC1 | 4.83 | 3.33 | 7.73E-04 |
| DDX47 | 4.83 | 3.04 | 7.75E-04 |
| MRGBP | 4.82 | 3.19 | 7.91E-04 |

| | | | |
|---|---|---|---|
| LRPPRC | 4.48 | 2.68 | 1.73E-03 |
| DERL2 | 4.47 | 2.73 | 1.74E-03 |
| INTS7 | 4.47 | 3.09 | 1.77E-03 |
| SDHB | 4.47 | 3.41 | 1.77E-03 |
| TARDBP | 4.47 | 2.62 | 1.78E-03 |
| BUD31 | 4.46 | 2.79 | 1.82E-03 |
| NUP88 | 4.45 | 2.87 | 1.85E-03 |
| PPCDC | 4.45 | 2.79 | 1.86E-03 |
| RPL11 | 4.44 | 3.19 | 1.87E-03 |
| DDX56 | 4.44 | 2.67 | 1.88E-03 |
| RBM33 | 4.44 | 2.73 | 1.91E-03 |
| EIF2B3 | 4.44 | 2.86 | 1.91E-03 |
| UMPS | 4.43 | 2.91 | 1.92E-03 |
| NHLRC2 | 4.43 | 3.03 | 1.93E-03 |
| NCAPG2 | 4.43 | 2.39 | 1.95E-03 |
| RBBP5 | 4.42 | 2.88 | 1.97E-03 |
| TARS2 | 4.42 | 3.07 | 1.99E-03 |
| NDUFAB1 | 4.41 | 2.84 | 2.03E-03 |
| ORAOV1 | 4.41 | 2.82 | 2.03E-03 |
| ATP6V0C | 4.4 | 3.38 | 2.06E-03 |
| AARS | 4.4 | 3.21 | 2.06E-03 |
| FAU | 4.4 | 2.63 | 2.07E-03 |
| EPRS | 4.38 | 2.66 | 2.14E-03 |
| RUVBL1 | 4.38 | 3.49 | 2.16E-03 |
| NAE1 | 4.38 | 2.81 | 2.17E-03 |
| COA7 | 4.36 | 2.72 | 2.26E-03 |
| MRPS31 | 4.36 | 3.06 | 2.26E-03 |
| EIF1AX | 4.36 | 2.8 | 2.28E-03 |
| GLRX5 | 4.35 | 2.97 | 2.29E-03 |
| PDRG1 | 4.35 | 2.92 | 2.30E-03 |
| YRDC | 4.35 | 2.86 | 2.30E-03 |
| FTSJ2 | 4.35 | 2.9 | 2.32E-03 |
| UBE2N | 4.32 | 2.88 | 2.47E-03 |
| MAK16 | 4.32 | 2.65 | 2.49E-03 |
| UBA1 | 4.32 | 2.72 | 2.50E-03 |
| MRPL43 | 4.32 | 2.78 | 2.51E-03 |
| SPATA5 | 4.3 | 2.99 | 2.60E-03 |
| KANSL3 | 4.3 | 2.54 | 2.61E-03 |
| MINOS1 | 4.3 | 2.99 | 2.64E-03 |
| MED30 | 4.29 | 2.98 | 2.65E-03 |
| POLR3E | 4.29 | 3.26 | 2.66E-03 |
| RPL4 | 4.29 | 2.79 | 2.70E-03 |
| MRPL15 | 4.28 | 2.81 | 2.71E-03 |
| PAFAH1B1 | 4.28 | 2.89 | 2.75E-03 |
| KIAA0391 | 4.27 | 2.74 | 2.76E-03 |
| MED19 | 4.27 | 2.57 | 2.77E-03 |
| PFDN1 | 4.27 | 2.9 | 2.79E-03 |

| | | | |
|---|---|---|---|
| CDC42 | 4.82 | 3.43 | 8.00E-04 |
| UBA2 | 4.82 | 2.67 | 8.01E-04 |
| PPP4C | 4.81 | 3.2 | 8.05E-04 |
| C10orf2 | 4.81 | 2.89 | 8.10E-04 |
| MRPL20 | 4.81 | 2.97 | 8.14E-04 |
| SMC3 | 4.81 | 3.15 | 8.14E-04 |
| PSMD14 | 4.8 | 2.71 | 8.22E-04 |
| RPS13 | 4.8 | 3.58 | 8.31E-04 |
| C21orf59 | 4.8 | 3.3 | 8.31E-04 |
| SART3 | 4.8 | 2.87 | 8.38E-04 |
| CPSF3L | 4.79 | 3.31 | 8.42E-04 |
| ERH | 4.78 | 3.48 | 8.68E-04 |
| HSPD1 | 4.78 | 3.27 | 8.84E-04 |
| GNB2L1 | 4.77 | 3.13 | 8.88E-04 |
| SYS1 | 4.76 | 3.1 | 9.13E-04 |
| RPLP0 | 4.76 | 3.24 | 9.15E-04 |
| CTCF | 4.76 | 3.45 | 9.15E-04 |
| RACGAP1 | 4.76 | 3.28 | 9.18E-04 |
| EIF2B3 | 4.76 | 2.94 | 9.20E-04 |
| MPHOSPH10 | 4.75 | 2.88 | 9.23E-04 |
| UBE2I | 4.75 | 3.43 | 9.23E-04 |
| TOMM22 | 4.75 | 2.86 | 9.23E-04 |
| SDHC | 4.75 | 2.96 | 9.40E-04 |
| SUPV3L1 | 4.74 | 3.13 | 9.55E-04 |
| MMS19 | 4.74 | 3.07 | 9.66E-04 |
| SUPT16H | 4.73 | 2.77 | 9.68E-04 |
| C1QBP | 4.73 | 2.72 | 9.77E-04 |
| RPS8 | 4.73 | 2.77 | 9.78E-04 |
| MRPS6 | 4.73 | 2.74 | 9.78E-04 |
| DDX11 | 4.73 | 2.76 | 9.84E-04 |
| TBCB | 4.72 | 2.92 | 9.91E-04 |
| PSMA6 | 4.72 | 2.74 | 1.00E-03 |
| RPL18 | 4.71 | 3.03 | 1.01E-03 |
| RPL7A | 4.71 | 2.67 | 1.01E-03 |
| CDIPT | 4.71 | 3.39 | 1.02E-03 |
| PGD | 4.71 | 3.06 | 1.02E-03 |
| PAM16 | 4.71 | 3.14 | 1.03E-03 |
| PDRG1 | 4.7 | 3.32 | 1.03E-03 |
| KRR1 | 4.7 | 3.53 | 1.03E-03 |
| RPL9 | 4.7 | 3.47 | 1.03E-03 |
| PRPF19 | 4.69 | 3.34 | 1.06E-03 |
| MRPL49 | 4.68 | 2.75 | 1.07E-03 |
| UTP15 | 4.68 | 2.75 | 1.09E-03 |
| PDCD11 | 4.67 | 3.34 | 1.10E-03 |
| EEF1G | 4.67 | 3.52 | 1.10E-03 |
| TRAPPC1 | 4.66 | 2.92 | 1.14E-03 |
| CSE1L | 4.65 | 2.9 | 1.17E-03 |

| | | | |
|---|---|---|---|
| CLNS1A | 4.27 | 2.8 | 2.79E-03 |
| MASTL | 4.27 | 2.96 | 2.81E-03 |
| PRPF38B | 4.26 | 2.99 | 2.83E-03 |
| TARS | 4.26 | 2.89 | 2.86E-03 |
| ARMC7 | 4.26 | 2.99 | 2.86E-03 |
| TRMT112 | 4.26 | 3.16 | 2.87E-03 |
| TIMELESS | 4.26 | 2.58 | 2.87E-03 |
| RPUSD3 | 4.25 | 2.93 | 2.91E-03 |
| TCP1 | 4.25 | 2.83 | 2.95E-03 |
| MRPS34 | 4.24 | 3.26 | 2.98E-03 |
| SNRPD1 | 4.24 | 2.56 | 2.98E-03 |
| DDX55 | 4.24 | 2.45 | 3.01E-03 |
| RNF4 | 4.23 | 2.31 | 3.05E-03 |
| GPN2 | 4.23 | 3.19 | 3.06E-03 |
| CCT4 | 4.22 | 3.01 | 3.12E-03 |
| MTG2 | 4.21 | 2.78 | 3.18E-03 |
| MRPL10 | 4.21 | 3.01 | 3.19E-03 |
| NKAP | 4.21 | 2.74 | 3.21E-03 |
| USP9X | 4.19 | 2.58 | 3.31E-03 |
| TOP2A | 4.19 | 2.38 | 3.32E-03 |
| RABIF | 4.18 | 2.29 | 3.39E-03 |
| COASY | 4.18 | 3.23 | 3.39E-03 |
| PCNA | 4.18 | 3.38 | 3.39E-03 |
| DDX10 | 4.18 | 2.77 | 3.41E-03 |
| PRPF3 | 4.18 | 2.48 | 3.42E-03 |
| CAPZB | 4.18 | 2.75 | 3.43E-03 |
| NUP43 | 4.18 | 2.86 | 3.44E-03 |
| CCNH | 4.18 | 3.06 | 3.44E-03 |
| CDK1 | 4.17 | 2.6 | 3.51E-03 |
| GARS | 4.17 | 2.57 | 3.52E-03 |
| TUBG1 | 4.17 | 2.7 | 3.53E-03 |
| ATL2 | 4.16 | 2.36 | 3.59E-03 |
| DHDDS | 4.16 | 2.59 | 3.61E-03 |
| TBC1D3F | 4.15 | 2.46 | 3.70E-03 |
| CDC37 | 4.14 | 3.03 | 3.74E-03 |
| RPL27 | 4.14 | 2.55 | 3.76E-03 |
| RPS8 | 4.14 | 2.99 | 3.79E-03 |
| RPL36 | 4.13 | 3.02 | 3.79E-03 |
| C9orf114 | 4.13 | 2.76 | 3.82E-03 |
| EXOSC1 | 4.13 | 2.6 | 3.87E-03 |
| RABGGTA | 4.13 | 2.73 | 3.89E-03 |
| PSTK | 4.12 | 2.84 | 3.91E-03 |
| RPAP1 | 4.12 | 2.92 | 3.91E-03 |
| NDUFA2 | 4.12 | 2.36 | 3.92E-03 |
| ERAL1 | 4.11 | 2.64 | 4.03E-03 |
| MED11 | 4.11 | 2.6 | 4.05E-03 |
| RPN2 | 4.11 | 3.26 | 4.05E-03 |

| | | | |
|---|---|---|---|
| PPWD1 | 4.65 | 3.25 | 1.18E-03 |
| RFC5 | 4.64 | 3.36 | 1.19E-03 |
| GRB2 | 4.64 | 2.63 | 1.20E-03 |
| HAUS7 | 4.64 | 3 | 1.20E-03 |
| MRPS25 | 4.63 | 2.99 | 1.21E-03 |
| EEF1A1 | 4.63 | 2.77 | 1.22E-03 |
| BNIP1 | 4.63 | 2.78 | 1.23E-03 |
| SNRPF | 4.63 | 3.35 | 1.23E-03 |
| ELL | 4.63 | 2.99 | 1.23E-03 |
| NUP133 | 4.61 | 2.69 | 1.29E-03 |
| XPO1 | 4.6 | 3.19 | 1.30E-03 |
| NFS1 | 4.6 | 3.49 | 1.30E-03 |
| PKMYT1 | 4.6 | 2.78 | 1.32E-03 |
| WDR43 | 4.6 | 2.97 | 1.32E-03 |
| SKP2 | 4.59 | 2.59 | 1.34E-03 |
| EIF2S1 | 4.59 | 2.92 | 1.34E-03 |
| ETF1 | 4.59 | 2.76 | 1.35E-03 |
| SNF8 | 4.59 | 2.87 | 1.35E-03 |
| WDR74 | 4.58 | 2.88 | 1.37E-03 |
| ACTL6A | 4.57 | 3.02 | 1.39E-03 |
| RPS20 | 4.57 | 3.04 | 1.41E-03 |
| RGPD6 | 4.57 | 2.58 | 1.41E-03 |
| RGPD5 | 4.57 | 2.58 | 1.41E-03 |
| SPC24 | 4.57 | 3.71 | 1.42E-03 |
| ECT2 | 4.56 | 3.02 | 1.42E-03 |
| PNPT1 | 4.56 | 3.03 | 1.42E-03 |
| UBA3 | 4.56 | 2.83 | 1.43E-03 |
| GOSR2 | 4.56 | 2.48 | 1.44E-03 |
| TXN | 4.56 | 3.42 | 1.44E-03 |
| SNUPN | 4.56 | 2.82 | 1.44E-03 |
| DENR | 4.55 | 2.74 | 1.45E-03 |
| ING3 | 4.55 | 2.75 | 1.46E-03 |
| WRB | 4.55 | 2.65 | 1.46E-03 |
| RPA1 | 4.54 | 2.92 | 1.48E-03 |
| DYNLL1 | 4.54 | 2.89 | 1.49E-03 |
| TSG101 | 4.54 | 2.65 | 1.50E-03 |
| PSMA3 | 4.53 | 2.89 | 1.52E-03 |
| MRPL53 | 4.53 | 2.94 | 1.52E-03 |
| CFAP20 | 4.53 | 2.53 | 1.53E-03 |
| SF3A3 | 4.53 | 2.85 | 1.53E-03 |
| PSMA7 | 4.52 | 2.93 | 1.55E-03 |
| ITGAV | 4.52 | 3.12 | 1.57E-03 |
| NUDT21 | 4.52 | 2.95 | 1.57E-03 |
| RRM1 | 4.52 | 3 | 1.57E-03 |
| AARS2 | 4.51 | 2.76 | 1.61E-03 |
| RPS29 | 4.51 | 2.69 | 1.61E-03 |
| CHMP4B | 4.51 | 3.26 | 1.62E-03 |

| | | | |
|---|---|---|---|
| WDR48 | 4.1 | 2.47 | 4.13E-03 |
| C12orf65 | 4.1 | 2.62 | 4.18E-03 |
| DNM1L | 4.08 | 2.92 | 4.40E-03 |
| DDX42 | 4.07 | 2.59 | 4.44E-03 |
| GGPS1 | 4.07 | 2.37 | 4.45E-03 |
| SRP14 | 4.07 | 2.74 | 4.51E-03 |
| MYBBP1A | 4.06 | 3.09 | 4.55E-03 |
| SKP2 | 4.06 | 2.36 | 4.55E-03 |
| KIF18A | 4.06 | 2.78 | 4.57E-03 |
| UTP3 | 4.06 | 2.67 | 4.58E-03 |
| IARS2 | 4.05 | 3.18 | 4.64E-03 |
| ENO1 | 4.05 | 2.68 | 4.71E-03 |
| DONSON | 4.04 | 2.6 | 4.80E-03 |
| HAUS7 | 4.03 | 3.07 | 4.94E-03 |
| CIAO1 | 4.03 | 2.89 | 4.95E-03 |
| POLG2 | 4.02 | 2.44 | 5.05E-03 |
| TANGO6 | 4.01 | 3.06 | 5.06E-03 |
| C10orf2 | 4.01 | 2.52 | 5.14E-03 |
| MRPL21 | 4 | 2.88 | 5.20E-03 |
| GNB2L1 | 4 | 3.32 | 5.25E-03 |
| MRPL2 | 4 | 2.31 | 5.27E-03 |
| ATP6V1G1 | 3.99 | 2.55 | 5.34E-03 |
| POLR3D | 3.98 | 2.65 | 5.40E-03 |
| USP5 | 3.98 | 2.86 | 5.41E-03 |
| RAD51D | 3.98 | 2.66 | 5.42E-03 |
| XPO1 | 3.97 | 2.77 | 5.43E-03 |
| RBBP4 | 3.97 | 2.67 | 5.46E-03 |
| PGS1 | 3.97 | 2.57 | 5.46E-03 |
| MRPS24 | 3.96 | 3.27 | 5.51E-03 |
| DHPS | 3.96 | 3 | 5.54E-03 |
| ADSL | 3.95 | 2.83 | 5.68E-03 |
| MRPL4 | 3.93 | 2.56 | 5.79E-03 |
| DDB1 | 3.93 | 3.1 | 5.79E-03 |
| SNRNP35 | 3.89 | 2.61 | 6.21E-03 |
| SEPHS2 | 3.88 | 3.12 | 6.26E-03 |
| ELP5 | 3.87 | 2.75 | 6.43E-03 |
| SMC1A | 3.86 | 2.8 | 6.46E-03 |
| HIST2H3A | 3.85 | 2.37 | 6.64E-03 |
| HIST2H3C | 3.85 | 2.37 | 6.64E-03 |
| MVD | 3.84 | 2.49 | 6.69E-03 |
| PRPF19 | 3.84 | 2.82 | 6.70E-03 |
| EEF1G | 3.84 | 2.87 | 6.76E-03 |
| MRTO4 | 3.83 | 2.82 | 6.80E-03 |
| HAUS5 | 3.83 | 2.98 | 6.82E-03 |
| ANKRD49 | 3.83 | 2.36 | 6.84E-03 |
| ZNF131 | 3.83 | 2.88 | 6.86E-03 |
| SHQ1 | 3.82 | 2.43 | 6.93E-03 |

| | | | |
|---|---|---|---|
| RPL19 | 4.51 | 2.82 | 1.62E-03 |
| NUP85 | 4.51 | 2.91 | 1.62E-03 |
| FAU | 4.5 | 3.03 | 1.63E-03 |
| EEF2KMT | 4.5 | 3.14 | 1.63E-03 |
| POP5 | 4.5 | 2.99 | 1.64E-03 |
| NAE1 | 4.49 | 2.95 | 1.67E-03 |
| NAA10 | 4.49 | 3.22 | 1.69E-03 |
| VRK1 | 4.48 | 2.9 | 1.70E-03 |
| DDX59 | 4.48 | 2.72 | 1.70E-03 |
| SNAPC1 | 4.48 | 2.78 | 1.72E-03 |
| SNRPE | 4.48 | 3.29 | 1.73E-03 |
| ALG2 | 4.48 | 3.01 | 1.73E-03 |
| ATP6V1A | 4.48 | 2.85 | 1.74E-03 |
| RPL3 | 4.47 | 2.85 | 1.74E-03 |
| NOL9 | 4.47 | 3.08 | 1.76E-03 |
| TSR2 | 4.47 | 3.09 | 1.76E-03 |
| RPS6 | 4.47 | 2.78 | 1.76E-03 |
| MASTL | 4.47 | 3.25 | 1.78E-03 |
| RAN | 4.46 | 2.62 | 1.78E-03 |
| NARS | 4.46 | 2.64 | 1.80E-03 |
| PSMA2 | 4.46 | 2.79 | 1.81E-03 |
| DHX33 | 4.46 | 3 | 1.81E-03 |
| DCLRE1B | 4.46 | 2.72 | 1.82E-03 |
| RPL5 | 4.45 | 3.18 | 1.82E-03 |
| RPS19 | 4.45 | 3.29 | 1.82E-03 |
| FAM96B | 4.45 | 2.85 | 1.84E-03 |
| ICE2 | 4.45 | 2.6 | 1.86E-03 |
| GTF2E1 | 4.44 | 3.06 | 1.89E-03 |
| TIMM13 | 4.44 | 2.44 | 1.89E-03 |
| LSM3 | 4.44 | 3.23 | 1.90E-03 |
| SRSF7 | 4.44 | 3.37 | 1.91E-03 |
| SEC13 | 4.43 | 2.62 | 1.93E-03 |
| MRPL22 | 4.43 | 2.7 | 1.93E-03 |
| SPCS3 | 4.43 | 3.28 | 1.94E-03 |
| DDX49 | 4.43 | 2.85 | 1.95E-03 |
| SRBD1 | 4.41 | 3.39 | 2.00E-03 |
| SRSF1 | 4.41 | 2.54 | 2.02E-03 |
| KIAA0391 | 4.4 | 2.95 | 2.05E-03 |
| POLE | 4.4 | 3.03 | 2.05E-03 |
| TIMM10 | 4.4 | 3.03 | 2.07E-03 |
| CDK7 | 4.39 | 3.6 | 2.08E-03 |
| TUBGCP5 | 4.39 | 2.54 | 2.08E-03 |
| WDR61 | 4.39 | 2.88 | 2.10E-03 |
| IMP4 | 4.39 | 2.74 | 2.10E-03 |
| NPIPA2 | 4.39 | 3.17 | 2.12E-03 |
| NPIPA1 | 4.39 | 3.17 | 2.12E-03 |
| NPIPA3 | 4.39 | 3.17 | 2.12E-03 |
| TBCB | 3.81 | 2.89 | 7.04E-03 |
| NUDC | 3.8 | 2.48 | 7.24E-03 |
| RPL12 | 3.79 | 2.77 | 7.27E-03 |
| NMT1 | 3.79 | 3.12 | 7.32E-03 |
| GNB2 | 3.79 | 2.44 | 7.36E-03 |
| GTPBP4 | 3.78 | 3.13 | 7.40E-03 |
| DDOST | 3.78 | 2.77 | 7.40E-03 |
| RARS2 | 3.78 | 2.78 | 7.44E-03 |
| RPP21 | 3.78 | 2.57 | 7.44E-03 |
| PLA2G10 | 3.76 | 2.51 | 7.74E-03 |
| NDUFA1 | 3.75 | 2.47 | 7.87E-03 |
| AK6 | 3.75 | 2.88 | 7.88E-03 |
| EP400 | 3.75 | 2.4 | 7.92E-03 |
| MRPL23 | 3.74 | 2.65 | 7.93E-03 |
| SF3A2 | 3.74 | 2.31 | 8.06E-03 |
| ACTR2 | 3.74 | 2.85 | 8.06E-03 |
| TELO2 | 3.73 | 2.31 | 8.21E-03 |
| SLC52A1 | 3.73 | 2.58 | 8.23E-03 |
| GTF2H1 | 3.71 | 2.69 | 8.40E-03 |
| MCM6 | 3.71 | 2.43 | 8.44E-03 |
| KPNB1 | 3.71 | 2.47 | 8.46E-03 |
| RRS1 | 3.7 | 2.84 | 8.57E-03 |
| IGBP1 | 3.69 | 2.64 | 8.85E-03 |
| TIMM13 | 3.69 | 2.44 | 8.86E-03 |
| UQCRFS1 | 3.68 | 2.33 | 8.94E-03 |
| NOP58 | 3.68 | 2.32 | 8.97E-03 |
| PAM16 | 3.68 | 2.51 | 9.01E-03 |
| MIS18A | 3.67 | 2.83 | 9.04E-03 |
| CUL1 | 3.67 | 2.61 | 9.04E-03 |
| TRIT1 | 3.67 | 2.68 | 9.16E-03 |
| MED6 | 3.66 | 2.47 | 9.28E-03 |
| PRMT5 | 3.65 | 2.78 | 9.44E-03 |
| URB1 | 3.64 | 2.79 | 9.66E-03 |
| IMP3 | 3.64 | 2.86 | 9.70E-03 |
| MRPL16 | 3.64 | 2.92 | 9.76E-03 |
| ATP6V1A | 3.63 | 2.82 | 9.81E-03 |
| BCLAF1 | 3.63 | 2.3 | 9.92E-03 |
| TRIAP1 | 3.62 | 2.65 | 9.98E-03 |
| HSD17B10 | 3.62 | 2.34 | 1.01E-02 |
| UQCC2 | 3.6 | 2.65 | 1.05E-02 |
| LANCL2 | 3.59 | 2.27 | 1.06E-02 |
| CSNK2B | 3.59 | 2.32 | 1.07E-02 |
| RMI1 | 3.58 | 2.57 | 1.08E-02 |
| NOL10 | 3.58 | 2.76 | 1.09E-02 |
| COX17 | 3.57 | 2.25 | 1.10E-02 |
| ALYREF | 3.57 | 2.26 | 1.12E-02 |
| PSMD7 | 3.56 | 2.64 | 1.12E-02 |

| | | | |
|---|---|---|---|
| KANSL3 | 4.38 | 2.39 | 2.17E-03 |
| NOP10 | 4.37 | 2.5 | 2.19E-03 |
| CDC37 | 4.37 | 2.98 | 2.20E-03 |
| RPL17 | 4.37 | 2.79 | 2.21E-03 |
| HUS1 | 4.37 | 2.8 | 2.21E-03 |
| RPL35A | 4.36 | 3.05 | 2.23E-03 |
| SMG1 | 4.36 | 2.73 | 2.24E-03 |
| MDN1 | 4.36 | 3.2 | 2.24E-03 |
| PGAM1 | 4.36 | 2.51 | 2.25E-03 |
| GTF2B | 4.36 | 3.02 | 2.28E-03 |
| XRN2 | 4.36 | 2.82 | 2.28E-03 |
| DAD1 | 4.35 | 3.57 | 2.30E-03 |
| ACTR10 | 4.35 | 2.59 | 2.30E-03 |
| PRMT5 | 4.35 | 3.26 | 2.30E-03 |
| RPL6 | 4.35 | 3.62 | 2.30E-03 |
| KPNB1 | 4.35 | 2.75 | 2.32E-03 |
| TRRAP | 4.35 | 2.62 | 2.34E-03 |
| ATP2A2 | 4.34 | 3.41 | 2.36E-03 |
| OXSM | 4.34 | 2.46 | 2.37E-03 |
| RRP12 | 4.34 | 2.78 | 2.38E-03 |
| RPS7 | 4.33 | 2.55 | 2.39E-03 |
| MED11 | 4.33 | 3.15 | 2.40E-03 |
| GNB1L | 4.33 | 2.64 | 2.40E-03 |
| FEN1 | 4.33 | 2.35 | 2.43E-03 |
| EIF1AD | 4.32 | 3.14 | 2.46E-03 |
| CARS | 4.32 | 3.11 | 2.46E-03 |
| POLE2 | 4.32 | 2.7 | 2.50E-03 |
| MCM7 | 4.32 | 2.85 | 2.51E-03 |
| RPS16 | 4.31 | 2.62 | 2.55E-03 |

| | | | |
|---|---|---|---|
| TBCA | 3.56 | 3.36 | 1.14E-02 |
| CPSF6 | 3.55 | 2.48 | 1.15E-02 |
| ATP5I | 3.55 | 2.68 | 1.15E-02 |
| HJURP | 3.55 | 2.25 | 1.15E-02 |
| ZRSR2 | 3.55 | 2.58 | 1.16E-02 |
| PPIL4 | 3.55 | 2.49 | 1.16E-02 |
| EIF2S1 | 3.54 | 2.58 | 1.17E-02 |
| HAUS1 | 3.54 | 2.84 | 1.18E-02 |
| NFRKB | 3.54 | 2.41 | 1.19E-02 |
| BIRC6 | 3.54 | 2.58 | 1.19E-02 |
| PPP1R12A | 3.53 | 2.36 | 1.19E-02 |
| DCTN3 | 3.53 | 2.55 | 1.20E-02 |
| EEF2KMT | 3.52 | 2.68 | 1.23E-02 |
| NAA10 | 3.52 | 2.91 | 1.24E-02 |
| OXSM | 3.51 | 2.19 | 1.25E-02 |
| RPS19 | 3.51 | 2.82 | 1.26E-02 |
| NOL6 | 3.5 | 2.56 | 1.28E-02 |
| SEPSECS | 3.49 | 2.83 | 1.30E-02 |
| CCDC51 | 3.48 | 2.46 | 1.32E-02 |
| RCC1 | 3.48 | 2.7 | 1.34E-02 |
| TTF2 | 3.48 | 2.34 | 1.35E-02 |
| SETD1A | 3.47 | 2.44 | 1.36E-02 |
| SNRNP27 | 3.47 | 2.22 | 1.36E-02 |
| MCM3 | 3.47 | 2.77 | 1.37E-02 |
| TXN | 3.47 | 2.34 | 1.37E-02 |
| RPS20 | 3.46 | 2.62 | 1.39E-02 |
| MCMBP | 3.46 | 2.21 | 1.40E-02 |
| DAD1 | 3.45 | 2.9 | 1.41E-02 |
| WDR43 | 3.45 | 2.81 | 1.43E-02 |
